# Unique functions for Notch4 in murine embryonic lymphangiogenesis

**DOI:** 10.1101/2021.02.19.431982

**Authors:** Ajit Muley, Minji Kim Uh, Glicella Salazar-De Simone, Bhairavi Swaminathan, Jennifer M. James, Aino Murtomaki, Seock Won Youn, Joseph D. McCarron, Chris Kitajewski, Maria Gnarra Buethe, Gloria Riitano, Yoh-suke Mukoyama, Jan Kitajewski, Carrie J. Shawber

**Author notes:** **Corresponding Author:** Carrie J. Shawber, Columbia University Medical Center, Physician and Surgeons, 16-440B, 630 W. 168th St., New York, NY 10032. These first authors contributed equally to this work (Ajit Muley, Minji Kim Uh).

## Abstract

In mice, embryonic dermal lymphatic development is well-understood and used to study gene functions in lymphangiogenesis. Notch signaling is an evolutionarily conserved pathway that modulates cell fate decisions, which has been shown to both inhibit and promote dermal lymphangiogenesis. Here, we demonstrate distinct roles for Notch4 signaling versus canonical Notch signaling in embryonic dermal lymphangiogenesis. Actively growing embryonic dermal lymphatics expressed NOTCH1, NOTCH4 and DLL4 which correlated with Notch activity. In lymphatic endothelial cells (LECs), DLL4 activation of Notch induced a subset of Notch effectors and lymphatic genes, which were distinctly regulated by Notch1 and Notch4 activation. Treatment of LECs with VEGF-A or VEGF-C upregulated *Dll4* transcripts, and differentially and temporally regulated the expression of *Notch1* and *Hes*/*Hey* genes. Mice nullizygous for *Notch4* had an increase in the closure of the lymphangiogenic fronts which correlated with reduced vessel caliber in the maturing lymphatic plexus at E14.5, and reduced branching at E16.5. Activation of Notch4 suppressed LEC migration in a wounding assay significantly more than Notch1, suggesting a dominant role for Notch4 in regulating LEC migration. Unlike *Notch4* nulls, inhibition of canonical Notch signaling by expressing a dominant negative form of MAML1 (DNMAML) in Prox1+ LECs led to increased lymphatic density consistent with an increased in LEC proliferation, described for the loss of LEC *Notch1*. Moreover, loss of *Notch4* did not affect LEC canonical Notch signaling. Thus, we propose that Notch4 signaling and canonical Notch signaling have distinct functions in the coordination of embryonic dermal lymphangiogenesis.

## Introduction

Lymphangiogenesis is the process by which new lymphatic vessels sprout off pre-existing vessels. Sprouting of new lymphatic vessels requires coordinated lymphatic endothelial cell (LEC) proliferation, directional migration, and cell-cell adhesion to form a properly patterned and functional network. In murine dorsal skin, lymphangiogenesis begins at embryonic day 12.5 (E12.5) at the side of the trunk and follows dermal blood vessel development to meet at the midline around E15.5 (Fig. S1a) [1]. Dermal lymphangiogenesis in mouse embryos is well characterized allowing for analysis of lymphatic endothelial signaling pathways, such as Notch.

The Notch family of signaling proteins consists of four cell surface receptors (NOTCH1-4) that are bound and activated by membrane-bound ligands of the Delta-like (Dll1, 4) and Jagged (Jag1, 2) families expressed on neighboring cells. Upon ligand activation, the extracellular domain of NOTCH is released, which induces conformational changes that expose two proteolytic cleavage sites (TACE and γ-secretase/presenilin) that in turn release of the intracellular cytoplasmic domain (NICD) from the cell surface [2]. In the canonical Notch signaling pathway, NICD transits to the nucleus, binds the transcriptional repressor RBPjκ, where it recruits an activation complex including Mastermind-like (MAML) and HDACs, and activates RBPjκ-dependent transcription of Notch effectors, such as those in the HES/Hey families. Notch also signals via a less well-understood non-canonical RBPjκ-independent pathway that has been suggested to not require nuclear localization of NICD [2].

During development of the blood vascular system, Notch signaling is essential for arterial endothelial specification, vascular smooth muscle cell differentiation and viability, and sprouting angiogenesis [3-5]. Studies of murine retinal angiogenesis have shown that VEGF-A, via activation of VEGFR2, upregulates DLL4 expression in the filopodia-extending tip cell located at the vascular front [6,4,7,8]. Dll4 signals to the neighboring Notch-expressing stalk cell, where Notch activation downregulates VEGFR2 and VEGFR3 expression and inhibits the tip cell phenotype. During retinal angiogenesis, inhibition of DLL4 or NOTCH1 leads to a hypersprouting phenotype characterized by an increase in tip cells at the expense of the stalk cells, increased VEGFR2 and VEGFR3 expression, and decreased vascular outgrowth [6,7,9]. Although it has been shown that VEGF-C induces DLL4 in cultured LECs [10,11], the mechanisms by which Notch regulates dermal lymphangiogenesis remain to be elucidated.

We previously demonstrated that NOTCH1 and NOTCH4 are expressed in the postnatal maturing dermal lymphatics [12]. Studies of postnatal lymphangiogenesis have shown that pharmacological inhibition or genetic manipulation of Dll4/Notch1 signaling can result in both increased and decreased lymphangiogenesis [13,11]. Neutralizing antibodies against NOTCH1 or DLL4 suppressed lymphangiogenesis in the postnatal mouse ear, tail dermis and a wounding model [13]. In contrast, an inhibitory soluble DLL4 extracellular domain fused to FC (Dll4FC) stimulated lymphangiogenesis in the postnatal mouse ear [11]. In embryonic dermal lymphangiogenesis, *Notch1* deletion in LECs did not affect lymphatic branching, but increased lymphatic vessel caliber which was proposed to be secondary to an increase in LEC proliferation and decreased LEC apoptosis [14]. More recently, it was shown that loss of one copy of *Dll4* was associated with reduced embryonic dermal lymphangiogenesis in mice [15], a phenotype opposite to that seen in retinal angiogenesis [7,8]. Additional studies are needed to clarify the differences in the lymphangiogenic phenotypes observed upon disruption of lymphatic endothelial Notch signaling.

Here, we examined the roles for Notch4 and canonical Notch signaling in embryonic dermal lymphangiogenesis. We demonstrated that NOTCH1, NOTCH4 and DLL4 are expressed, and Notch signaling active in embryonic dermal lymphatic endothelium. VEGF-A and VEGF-C signaling differentially regulated Notch pathway gene expression and activity in cultured LECs. Mice nullizygous for *Notch4* displayed an embryonic dermal lymphangiogenic phenotype characterized by increased LEC migration and reduced branching. In contrast, inhibition of canonical Notch signaling increased lymphatic vascular density consistent with an increased in LEC proliferation. Together, these data demonstrate that dermal lymphangiogenesis is dynamically regulated by Notch and requires both NOTCH1 and NOTCH4 functions, as well as canonical and non-canonical Notch signaling.

## Materials and Methods

### Cell culture/Constructs

HeLa cells were maintained in 10% FBS DMEM. Human umbilical vein endothelial cells (HUVEC) were isolated as previously described and maintained in EGM2 (Lonza) [16,17]. Neonatal human dermal lymphatic endothelial cells (HdLECs) were either purchased (Promocell) or isolated as previously described [18], and maintained on fibronectin-coated plates in EGM2-MV2 (Lonza; complete medium) supplemented with 10 ng/mL VEGF-C (R&D). To activate Notch signaling, HdLECs were lentivirally infected [19] using pCCL.pkg.wpre vector encoding N1IC, N4/Int-3 or GFP. N1IC encodes the constitutively active cytoplasmic domain of NOTCH1. N4/Int-3 encodes an activated *Notch4* allele generated by MMTV insertion [20]. Protein expression was confirmed by quantitative (q)RT-PCR and Western analyses of samples collected post-infection.

### HdLEC assays

VEGF treatment of HdLECs: Confluent HdLEC monolayers were starved overnight in 1% FBS in EBM2 (Lonza) or in human endothelial SFM (Fisher Scientific) followed by either 1 hour or 5 hours in EBM2/SFM containing 100ng/mL VEGF-A (R&D), 100ng/mL VEGF-C (R&D), or 500ng/mL VEGF-C C156S (R&D) prior to RNA isolation. Assays were performed at least 3 times. For detection of AKT and ERK activity, HdLECs were serum starved overnight in SFM containing 1% FBS and 0.1% BSA, followed by 5 hours in SFM alone. Cells were then switched to SFM containing 0.1%BSA and either 100ng/mL VEGF-A or 100ng/mL VEGF-C for 20 minutes prior to fixation with cold 4% PFA. Assays were performed in duplicate for two different lentiviral transductions.

#### Migration Assay

HDLECs were seeded in triplicate on a fibronectin-coated (Thermofisher) 12 well plate in complete medium. The following day (0-hour time-point), a scratch through the confluent monolayer was made across each well using a yellow pipet tip, and medium was changed to EBM2 containing 100ng/ml VEGF-C. For migration assays with mitomycin C, confluent monolayers were pretreated with 10ug/mL mitomycin-C (Sigma) for 45 minutes prior to scratching. Cells were maintained in EBM2 containing 100ng/ml VEGF-C and 0.1 μg/mL mitomycin-C while migration was assessed. Growth into the scratch was documented at 0, 4, 8, 12, and 25 hours with a Zeiss Axiovert 40 CSL inverted microscope or at 0, 4, 8 and 24 hours using an Olympus IX83 microscope. Cell migration rate was determined using imageJ software [21], and calculated as the percentage of cell-free area at different time-points relative to the initial wound area. Assays were performed at least 2-3 times for two independent lentivirally-generated HDLEC populations.

### Co-culture Notch Reporter Assay

HDLECs (90% confluent) were lipofected (Lipofectamine 2000; Invitrogen) with the Notch reporter plasmid pGL3.11CSL [12] containing 11 repeats of Notch/CSL (RBPjκ) cis-elements, and phRL-SV40 *renilla* (Promega) to normalize lipofection efficiency. HeLa cells were lipofected with pCR3 plasmids encoding either DLL4-FLAG or JAG1-FLAG with empty vector serving as a control. 24 hours after lipofection, HeLa and HdLECs were co-cultured together at a 1:1 ratio on fibronectin-coated plates in EGM2. 24 hours after co-culture, a luciferase reporter assay was performed using the Dual Luciferase Reporter Assay System (Promega) and a TD20/20 luminometer (Turner Designs). Luciferase values were normalized to Renilla values. Each condition was performed in triplicate, 4 times.

### DLL4-ligand activation assay and mRNA sequencing

Tethered Ligand Assay: The recombinant extracellular domain of the Notch ligand hDLL4FC (Sino Biologicals Inc.) or IgG-FC (Sino Biologicals Inc.) were coated onto a 24-well plate (Corning) on a fibronectin matrix (Sigma). Following an overnight incubation at 4°C, primary ECs (at 80% confluency) were trypsinized and seeded onto the coated plates and incubated at 37°C with 5% CO_2_ for 6 hours. Experiment was performed in triplicate.

RNA was isolated using the RNEasy Mini Kit (Qiagen), quantity and integrity measured using a Bio-analyzer (Agilent TapeStation 4200, UIC Genome Research core) prior to RNA sequencing. TLA HdLEC samples were sequenced at a ∼30 million paired-end (PE) read depth with 150-base fragments by Novogene (https://en.novogene.com/). Raw reads from in-vitro screens were mapped to the Human database (ENSEMBL/GRCh38) using STAR (version 2.5.0a) and processed with Samtools (version 1.4.1). The counts obtained by FeatureCounts (version 1.5.2) were analyzed by DESeq2 (version 1.18.1) to identify differentially expressed genes. The datasets generated during and analyzed during the current study are available in the NCBI Gene Expression Omnibus repository at https://www.ncbi.nlm.nih.gov/geo (Accession number GSE183631).

### Gene Expression Analyses

RNA was isolated using the RNEasy Mini Kit (Qiagen) and reverse transcribed using the VersoTM cDNA Synthesis Kit (Thermo Fisher) or First Strand Synthesis Kit (Invitrogen). qRT-PCR was performed in triplicate for each gene (Table S1), using ABsoluteTM Blue QPCR SYBR Green Master Mix (Thermo Fisher) or Sybr Green Master Mix (ABI) and 7300 Real-Time PCR System (Applied Biosystems) or CFX96 PCR Cycler (Biorad). Gene specific qRT-PCR standards were used to determine transcript levels and normalized to *β-actin* expression [12]. Analyses were set up in triplicate and performed at least 3 times.

For the validation of mRNA sequencing data, qPCR was done using SYBR Green master mix (Applied Biosystems, Cat.4385612) and primers specific to genes of interest (Table S2). The mean cycle threshold (Ct) values from the triplicate run for each sample were analyzed using *β-actin* as the reference gene. ΔΔCt method [22] was used to calculate the relative expression using the following steps: 1) Normalization to reference gene: ΔCt_GOI_ = Ct_GOI_ – Ct_BA_. 2) Relative expression between conditions: ΔΔCt_GOI_ = ΔCt_EXP_ - ΔCt_CNT_. The analysis was done using Microsoft Excel and Prism.

### Western blotting

NOTCH4 expression was determined by Western blot. Fresh E14.5 hindlimbs were lysed in RIPA buffer (Invitrogen) containing protease and phosphatase inhibitors (Thermofisher) on ice and protein concentration determined by BCA protein assay kit (Pierce). Equal amounts of protein were separated on an 8% SDS-PAGE gel and transferred to a nitrocellulose membrane. The membranes were blocked with 3% nonfat milk and 3% bovine serum albumin (Jackson ImmunoResearch) in Tris-Buffered Saline Tween-20 and probed with antibodies against the cytoplasmic domain of NOTCH4 [23] and β-ACTIN (Abclonal). Horse radish peroxidase-conjugated secondary antibodies (Life Technologies) were used, detected with SuperSignal West Pico Chemiluminescent Substrate (Pierce) and images captured with Biorad Chemdoc MP.

### Mouse studies

*Notch4* nullizygous (*N4*^*-/-*^) [24], *Prox1CreER*^*T2*^ [25], and *DNMAML*^*fl/fl*^ [26], *CBF:H2B-Venus* (*NVR* reporter purchased from Jax Labs) [27] and *Prox1-tdTomato* lymphatic reporter (*ProxTom*); [28] mice were used for these studies. Studies were performed in mice in a mixed background, as well as a pure C57BL6j background. For studies using *Prox1CreER*^*T2*^, tamoxifen in corn oil was administered via oral gavage (10 mg/40 g) at E12.5. 3 or more independent litters were assessed for each analysis. Number of embryos analyzed is presented in the figure legends.

### Immunohistochemistry & Imaging

E14.5 and E16.5 dorsal skin was dissected, fixed for 2 hours in 4% PFA and then immunostaining initiated. Alternatively, embryos were incubated overnight in 4% PFA and then stored in 1 x PBS at 4**°**C. E14.5 tissues were incubated for 2 hours at room temperature in blocking buffer (10% donkey serum, 0.3% Triton X-100, 1 x PBS), incubated in primary antibody (Table S3) diluted in blocking buffer overnight at 4**°**C, and then incubated with the appropriate Alexa-Fluor secondary antibodies (Invitrogen) diluted in blocking buffer overnight at 4**°**C. E16.5 dermal tissues were washed in 1 x PBS containing 0.2% Triton X-100 and 20% DMSO for 4 hours at room temperature and immunohistochemistry performed as described in Cha et al 2016 [29]. For immunostaining of sections, 5-micron sections were stained as previously described [30]. Tissue was mounted using Vectashield with and without DAPI (Vector Laboratories). Images were captured using a Nikon SMZ-U Zoom 1:10 microscope and Nikon 4500 digital camera, Nikon ECLIPSE E800 microscope and NIS Elements software, Nikon DXM 1200 digital camera, and Image ProPlus v.4.01 software, a Zeiss Axioskop2 Plus and Zeiss AxioCam MRc camera with Zeiss Zen software, or an Olympus IX83 Inverted System Microscope and Olympus cellSens software. Confocal microscopy was performed with a Zeiss LSM 510 META Confocal Microscope and the LSM software.

For cell immunochemistry, cells were fixed in 4% PFA on ice for 15 minutes, permeabilized and blocked with 0.1% Triton X-100, 2% BSA, 3% donkey serum in 1 x PBS for 1 hour at room temperature. Cells were then incubated overnight with primary antibody at 4°C followed by an incubation with Alexa Fluor-conjugated anti-donkey-secondary antibodies (Invitrogen) for 1 hour at room temperature. Slides were washed in 1 x PBS and mounted with Vectashield with DAPI mounting media. Images were captured with an Olympus IX83 Inverted System Microscope and Olympus CellSens software.

Images were analyzed with ImageJ or Adobe Photoshop. Tiled 10x images were used to quantify lymphatic and blood vascular density, distance between migration fronts, fronts per unit length, and branch-point per unit length. 20x images were used to determine lymphatic vessel caliber, distance to first branch-point, number of Prox1+ cells per field, and sprout morphology. Vascular density was determined as positive signal area normalized by total area. Distance between migrating fronts was determined as the mean distance between the 2 lymphatic fronts measured at multiple points (≥3) [1]. For analysis of *Notch4* mutants, *Notch4*^+/-^ embryos were used to normalize between litters, as they were present in all litters analyzed and the distances between migration fronts were not statistical different than WT embryos. Sprouting fronts, defined as the sprouts that reside at the leading edge of the migrating front, per unit length was determined as the number of sprouting fronts at the leading edge of the migration front normalized to the vertical length (posterior-anterior; Fig. S1a). Length of the sprout was determined as length from tip of sprout at lymphangiogenic front to the first branch-point. Lymphatic vessel caliber was determined by measuring the width of lymphatic vessels in the maturing lymphatic plexus and adjacent to the first branch-point away from the migrating front. Sprout morphology at the lymphangiogenic front was determined by counting total number of blunt-ended sprouts (rounded, lacking multiple filopodia) and spiky-ended sprouts (elongated with multiple filopodia) normalized to the total number of sprouts assessed. Branch-points per unit length in maturing lymphatic plexus was determined as the number of branch-points per field normalized to the total length of lymphatic vessels per field. To measure Prox1+ LEC number, Prox1+/LYVE1+ LECs were scored and mean number per field determined. To determine the significance between control and one experimental group, a two-tailed student’s T-Test was used. For analyses of more than two groups, one-way analysis of variance (ANOVA) was used to determine significance by unpaired T-Test. For analyses of multiple conditions and cell populations, two-way ANOVA was used and Dunnett’s multiple comparison test performed to determine significance between groups. A p<0.05 was considered significant.

### Lymphangiography

Lymphangiographies were performed as described on E17.5 embryos [15]. Briefly, 2 µL of 0.4% Trypan blue solution (Sigma) was injected into the dermis in periorbital region with a 36G beveled needle attached to a Nano l syringe (WPI). The embryos were imaged using AMSCOPE stereomicroscope (AMSCOPE) with camera attachment 1 minute after injection.

## Results

### Embryonic dermal lymphatics expressed NOTCH1, NOTCH4 and the Notch ligand, DLL4

Cultured human LECs, HdLECs, express NOTCH1-4 and the Notch ligands, DLL4 and JAGGED1 (JAG1) [18], while NOTCH1 and NOTCH4 are expressed in the P4 murine dermal lymphatic vessels [12]. Notch signaling has been shown to be active in the E15.5 dermal lymphatics [14], but it is not known which Notch proteins and ligands are expressed in dermal LECs at this time. To study the role of Notch signaling in embryonic dermal lymphangiogenesis, we determined the expression of NOTCH1 and NOTCH4, and the angiogenic Notch ligands, DLL4 and JAG1, as well as Notch activity in E14.5 dorsal skin. This time-point is characterized by the presence of two LYVE1+ lymphatic fronts migrating towards the midline which precedes a maturing lymphatic plexus (Fig. S1a). At this time-point, two CD31+ angiogenic fronts have fused at the midline to form a connected blood capillary network.

At E14.5, DLL4 was expressed in both in the developing lymphatics and blood vessels of the dermis (Fig. 1a). Unlike the retina, where DLL4 expression is restricted to 1-2 tip cells at the angiogenic front [6-8], high DLL4 expression was observed in multiple LECs within the sprouts at the lymphangiogenic front. DLL4 was also expressed in the arterial vessels and blood capillary network, with strongest expression observed in the large arteries, consistent with its expression in the vasculature of the intestinal villi [10]. Unlike DLL4, JAG1 was not expressed in dermal lymphatics at E14.5 (Fig. 1a). JAG1 expression was limited to the blood vasculature, where the highest expression was observed in the larger caliber arteries in a pattern consistent with vascular smooth muscle cell and endothelial cells. Staining for NOTCH4 and NOTCH1 demonstrated that they were both expressed through-out the endothelium of the sprouts at the lymphatic front which overlapped with DLL4 (Fig. 1b, c). Analysis of E14.5 dermal cross-section confirmed the LYVE1+ dermal lymphatic endothelium expressed both NOTCH1 and NOTCH4 (Fig. S1b, c). Outside of the lymphatics, NOTCH1 expression was observed in the epidermis and blood endothelium, while NOTCH4 was expressed in the epidermis and a subset of LYVE1+ macrophages (Fig. 1b, c, Fig. S1b, c).

**Fig. 1.**
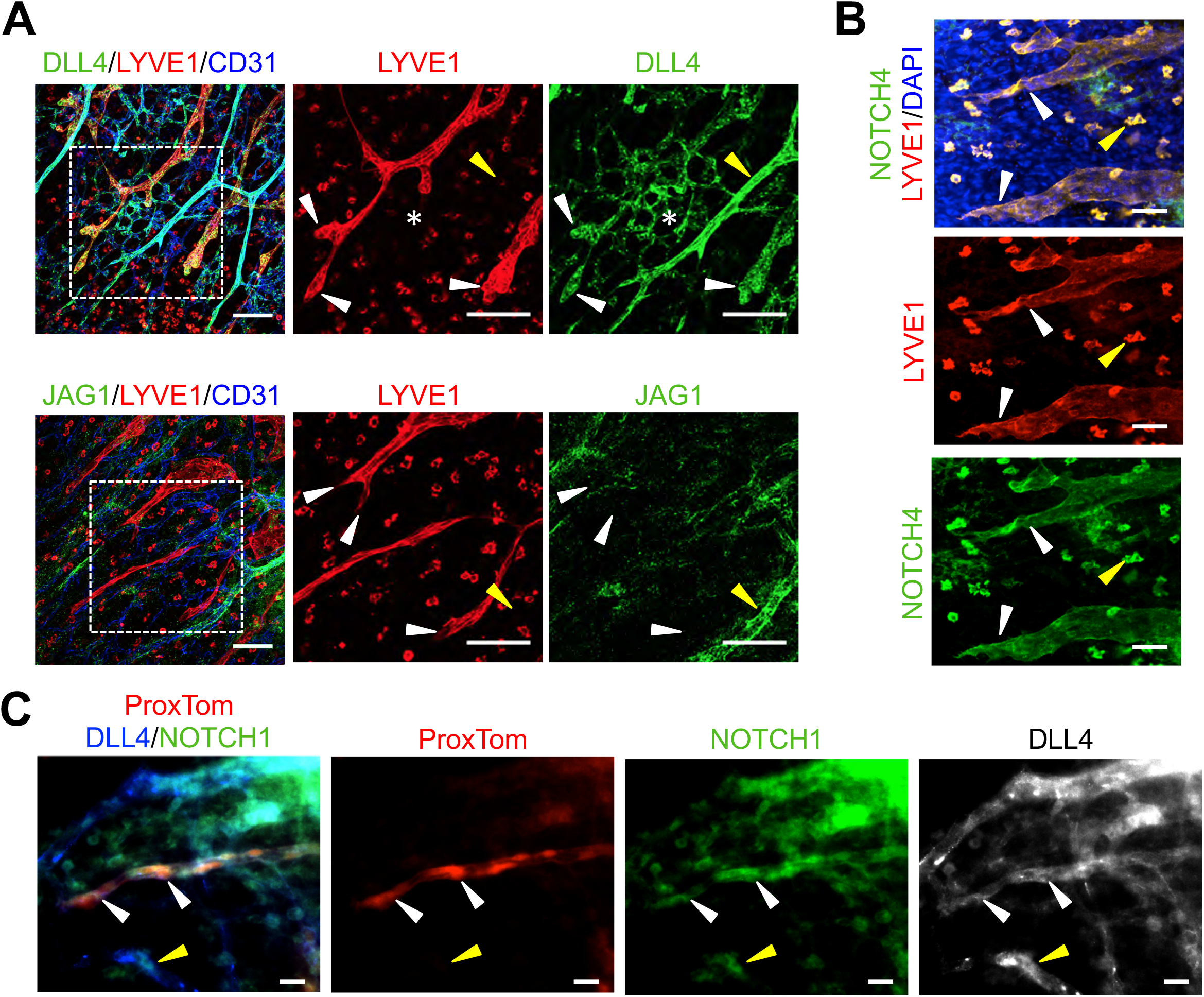
Embryonic dermal lymphatics expressed NOTCH1, NOTCH4 and DLL4. **a**) E14.5 wild-type skin wholemounts stained for LYVE1, CD31 and DLL4 or JAG1. Higher magnification of boxed areas presented to the right. White arrowheads mark sprouts at the lymphangiogenic front. White asterisk marks the blood vascular plexus. Yellow arrowhead marks an artery. Scale bars, 100µm. **b**) E14.5 wild-type skin wholemount stained for LYVE1 and NOTCH4. White arrowheads mark lymphatics at the front. Yellow arrowhead marks a NOTCH4+ macrophage. Scale bars, 50µm. **c**) E14.5 *ProxTom* skin wholemount stained for DLL4 and NOTCH1. White arrowheads mark lymphatic sprout at the front. Yellow arrowhead mark NOTCH1+ blood vessel. Scale bars, 20µm.

To determine where Notch is actively signaling during dermal lymphangiogenesis, the dermal lymphatics in E14.5 embryos carrying alleles for the Prox1-Tomato (*ProxTom*) LEC reporter [28] and the Notch Venous Reporter (*NVR*) [27] were assessed. Notch activity was observed throughout the lymphatic vascular plexus at both the lymphangiogenic front, defined as the LECS that make up the sprout from tip to the first branch point, and the mature plexus where the vessels have begun to remodel (Fig. 2a). At the lymphangiogenic front, Notch activity was often observed in several LECs located at the tip-cell positions in spiky-ended sprouts with filopodia (Fig. 2b), consistent with the broad expression of NOTCH1, NOTCH4 and DLL4 at the front (Fig. 1). Notch activity was also observed in blunt-ended sprouts. In the mature lymphatic plexus, Notch activity was observed at branch-points (Fig. 2a, c). Taken together the expression data suggest DLL4 signaling via NOTCH1 and/or NOTCH4 has a role in regulating dermal lymphangiogenic growth and maturation.

**Fig. 2.**
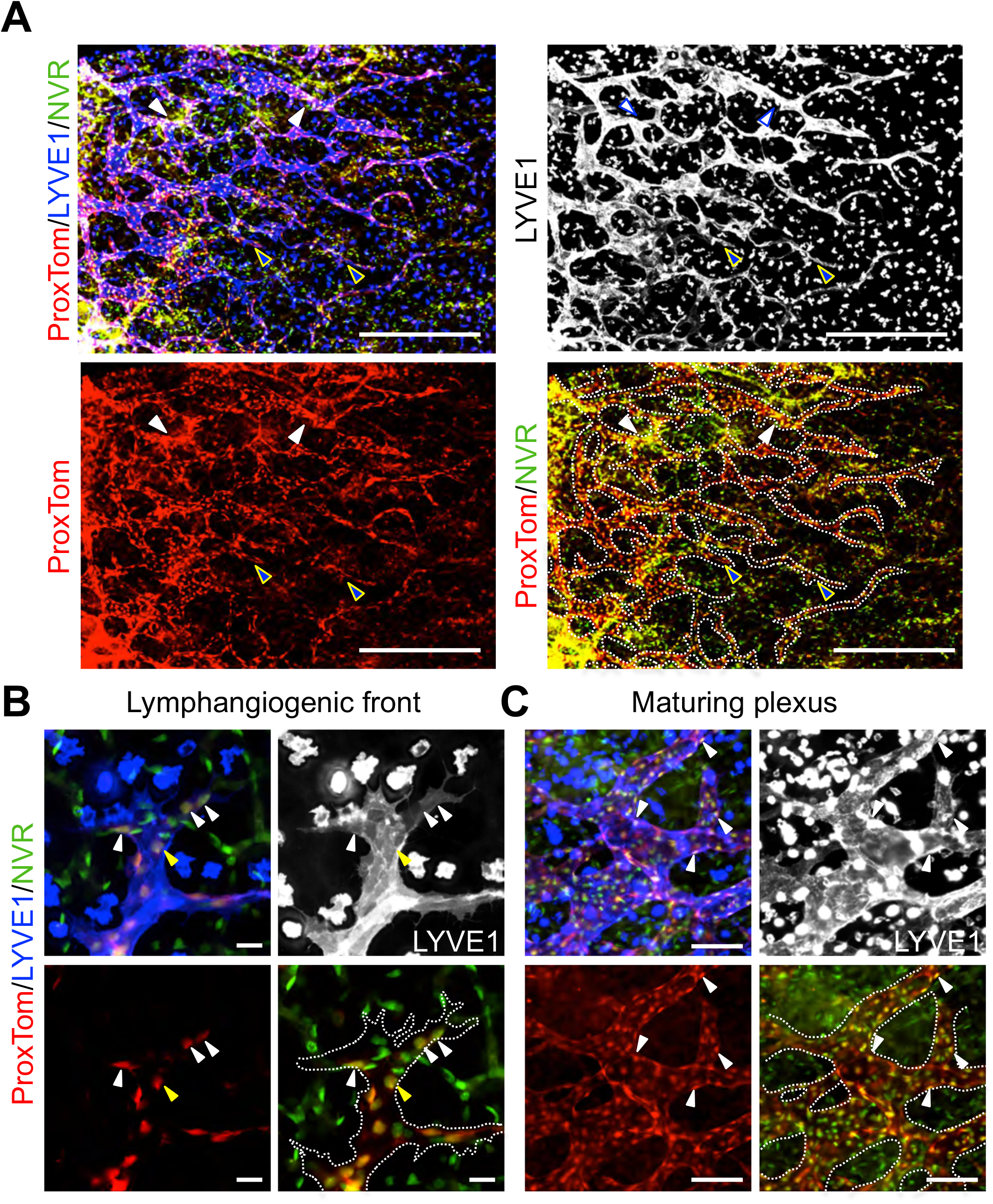
Notch activation observed throughout the embryonic dermal lymphatic vascular plexus. E14.5 *ProxTOM;NVR* skin wholemounts stained for LYVE1. **a**) Low magnification image demonstrating Notch activity throughout the developing lymphatic plexus. Blue arrowheads mark sprouts at the lymphangiogenic front with Notch activity. White arrowheads mark regions of high Notch signaling in the maturing plexus. Scale bars, 500µm. **b**) High magnification of spiky-ended lymphatic sprout. White arrowheads mark tip-cells with Notch activity. Yellow arrowheads mark stalk cells with Notch activity. **c**) High magnification of the maturing plexus. White arrowheads mark LECs with Notch activity. **b, c**) Scale bars, 100µm.

### Profiling of DLL4/Notch signaling in human dermal lymphatic endothelial cells

Expression studies suggested that DLL4 is the major ligand for Notch signaling in the lymphatic endothelium during dermal lymphangiogenesis. To determine if DLL4 or JAG1 could activate Notch in LECs, co-culture assays were performed in which endogenous Notch activation was determined using a Notch-response CSL luciferase reporter [12]. HeLa cells were engineered to express DLL4, JAG1 or both (Fig. S2a), and then seeded at a 1:1 ratio with HdLECs containing a CSL-luciferase reporter. DLL4-expressing HeLa cells upregulated Notch signaling nearly 5-fold over co-cultures using parental HeLa cells, while JAG1 only modestly increased Notch signaling in HDLECs (Fig. S2b). Co-culture with HeLa cells co-expressing DLL4 and JAG1 induced Notch signaling similar to the co-cultures with DLL4 alone, suggesting that JAG1 did not interfere with DLL4 signaling. Together with the expression studies, these data suggest that DLL4 functions as a ligand for LEC NOTCH.

To further assess DLL4/Notch signaling, HdLECs were seeded on DLL4FC-coated or FC-coated (control) plates. After 6 hours, RNA was collected and mRNA sequencing performed. Relative to the HdLECs seeded on FC-coated plates, HdLECs seeded on DLL4FC significantly altered the expression of 675 genes with a padj <0.05 (Fig. 3a, Table S4, S5). 69 genes were induced 1.2-fold, while 68 genes were suppressed 1.2-fold. Analysis of the top 30 upregulated genes revealed DLL4/Notch signaling upregulated the expression of known direct effectors of Notch signaling, *Hey1, Hes4, Dll4*, and *Hes1*, as well as key lymphangiogenic genes, *Ackr3, Cxcr4, Ccl2, EphrinB2, Gja4* (Cx37), *Gja1* (Cx43), and *Sema3g* (Fig. 3b) [31-38]. DLL4/Notch signaling also downregulated lymphangiogenic genes, including *Apln* and *Adm* (Fig. 3c) [39,35]. Further analysis of the 675 altered genes demonstrated that DLL4/Notch signaling both upregulated and downregulated genes of the Notch pathway (Fig. S3a) and lymphangiogenesis (Fig. S3b). GO: Biological Pathway (BP) analysis indicates that LEC DLL4/Notch signaling induces genes responsible for pattern specification, neurogenesis and chemotaxis (Fig. 3d).

**Fig. 3.**
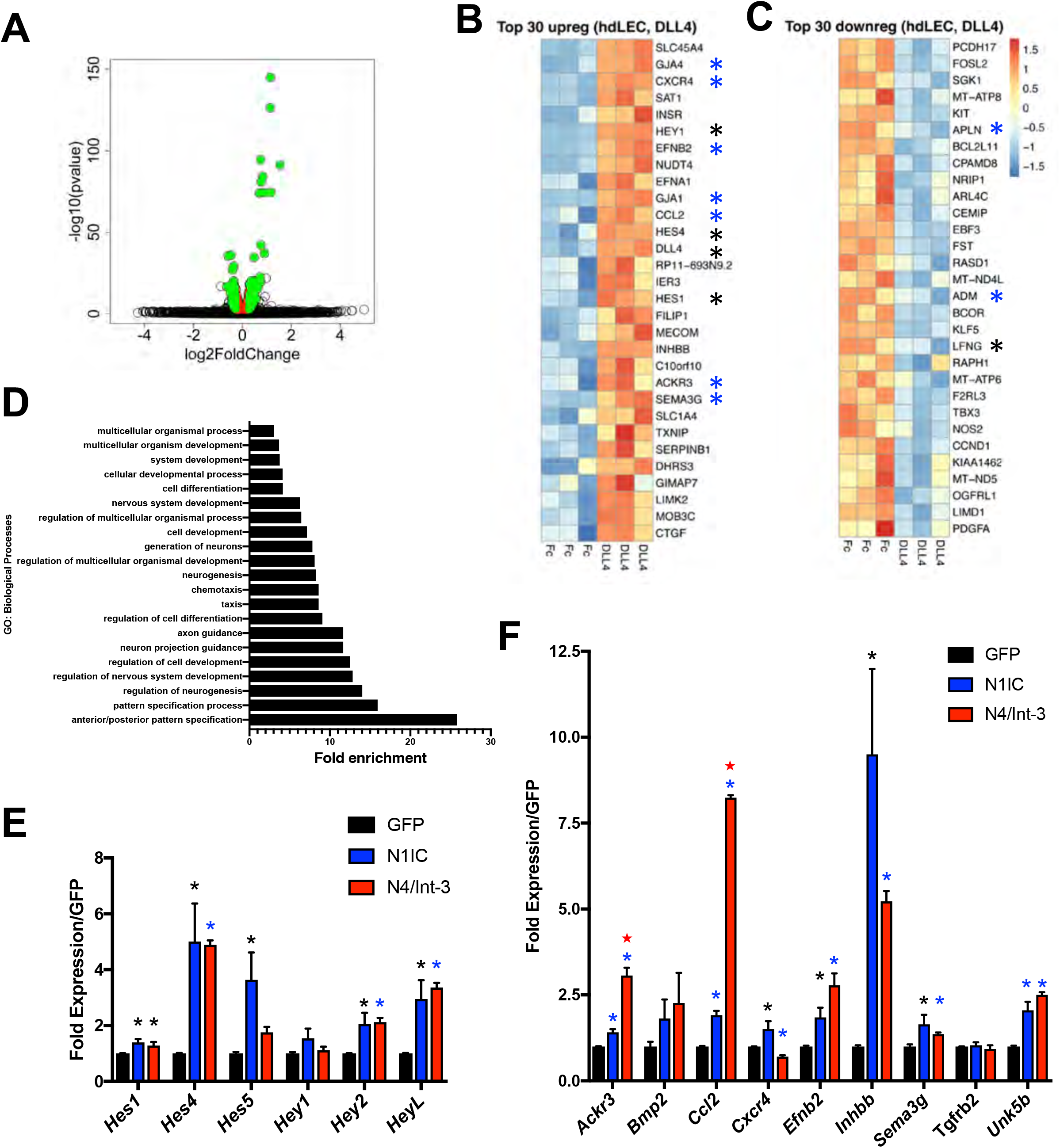
DLL4/Notch signaling regulated Notch and lymphangiogenic genes via Notch1 and Notch4. HdLECs were seeded on either DLL4FC- or FC-coated plates, and RNA isolated after 6 hours, followed by mRNA sequencing. Experiment was performed in triplicate. **a)** Volcano plot of genes downregulated and upregulated by DLL4FC relative to FC controls. **b**) Top 30 genes upregulated and **c**) downregulated by DLL4FC in HdLECs. **b, c**) * mark Notch pathway and 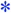 lymphangiogenic genes. **d)** Top GO pathways for biological processes for DLL4 induced genes. **e)** qRT-PCR for direct targets of Notch signaling and **f)** lymphangiogenic genes significantly induced in the DLL4-HdLEC assay in HdLECs expressing GFP, N1IC or N4/Int-3. Data presented for two independent transductions done in duplicate and gene expression determined by delta CT method and presented relative GFP controls ± s.e.m. One-way ANOVA performed and significance determined by unpaired T-test. **e, f**) * p<0.05 or 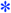 N4/Int-3 or N1IC relative to GFP controls. 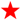 p<0.0003 N4/Int-3 relative to N1IC.

To assess if DLL4-induced genes were downstream of Notch1 or Notch4 activation, HdLECs were generated to express activated forms of NOTCH1 (N1IC) or NOTCH4 (N4/Int-3). Quantitative RT-PCR was performed for Notch effectors of the *Hes* and *Hey* gene families, as well as lymphangiogenic genes. Of the Notch effectors assessed, N1IC and N4/Int-3 both induced the expression of *Hes1, Hes4, Hey2* and *HeyL*, with the strongest induction observed for *Hes4* and *HeyL* (Fig. 3e). In contrast, only N1IC induced the expression of *Hes5*. Notch1 and Notch4 activation significantly induced the expression of the majority of lymphangiogenic genes assessed, except for *Cxcr4, Bmp2*, and *Tgfrb2* (Fig. 3F). Similarly, both suppressed the expression of *Prox1, Podoplanin* and *Lyve1* (Fig. S3b, S4b). *Cxcr4* was significantly induced by N1IC, while N4/Int-3 suppressed its expression. Neither *Tgfrb2* and *Bmp2* were induced. While *Ackr3* and *Ccl2* were significantly induced by N1IC relative to the GFP controls, N4/Int-3 was a much stronger inducer of both these genes (p<0.0003, p<0.0001 N4/Int-3 vs. N1IC, respectively). Together these data support overlapping and distinct downstream signaling for Notch1 and Notch4 in LECs.

### VEGF-C induced Dll4 expression and Notch activation in HdLECs

During sprouting angiogenesis, VEGF-A/VEGFR-2 signaling upregulates DLL4 in blood endothelial tip cells to activate Notch signaling in the adjacent stalk cell [6-8]. As we observed DLL4 expression and Notch activity in LECs of the sprouts located at the lymphangiogenic front, we determined the effect of VEGF-A and VEGF-C on Notch genes, ligands and effectors in HdLECs. Serum starved HdLECs were treated with either VEGF-A, VEGF-C or VEGF-C^C156S^. In HdLECs, VEGF-A binds and activates VEGFR2, VEGF-C^C156S^ binds and activates VEGFR3, and VEGF-C activates both VEGFR2 and VEGFR3 [40]. After 1 hour, VEGF-A and VEGF-C significantly induced *Dll4* transcripts, which correlated with an increase in *Hey1, Hey2* and *Hes1* transcripts (Fig. 4a, b). VEGF-C or VEGF-C^C156S^ both induced *Dll4* expression, as well as the Notch effector, *Hes1* (Fig. 4a, b). *Notch1* was modestly induced by VEGF-A only, while *Notch4* expression was unaffected (Fig.4a).

**Fig. 4.**
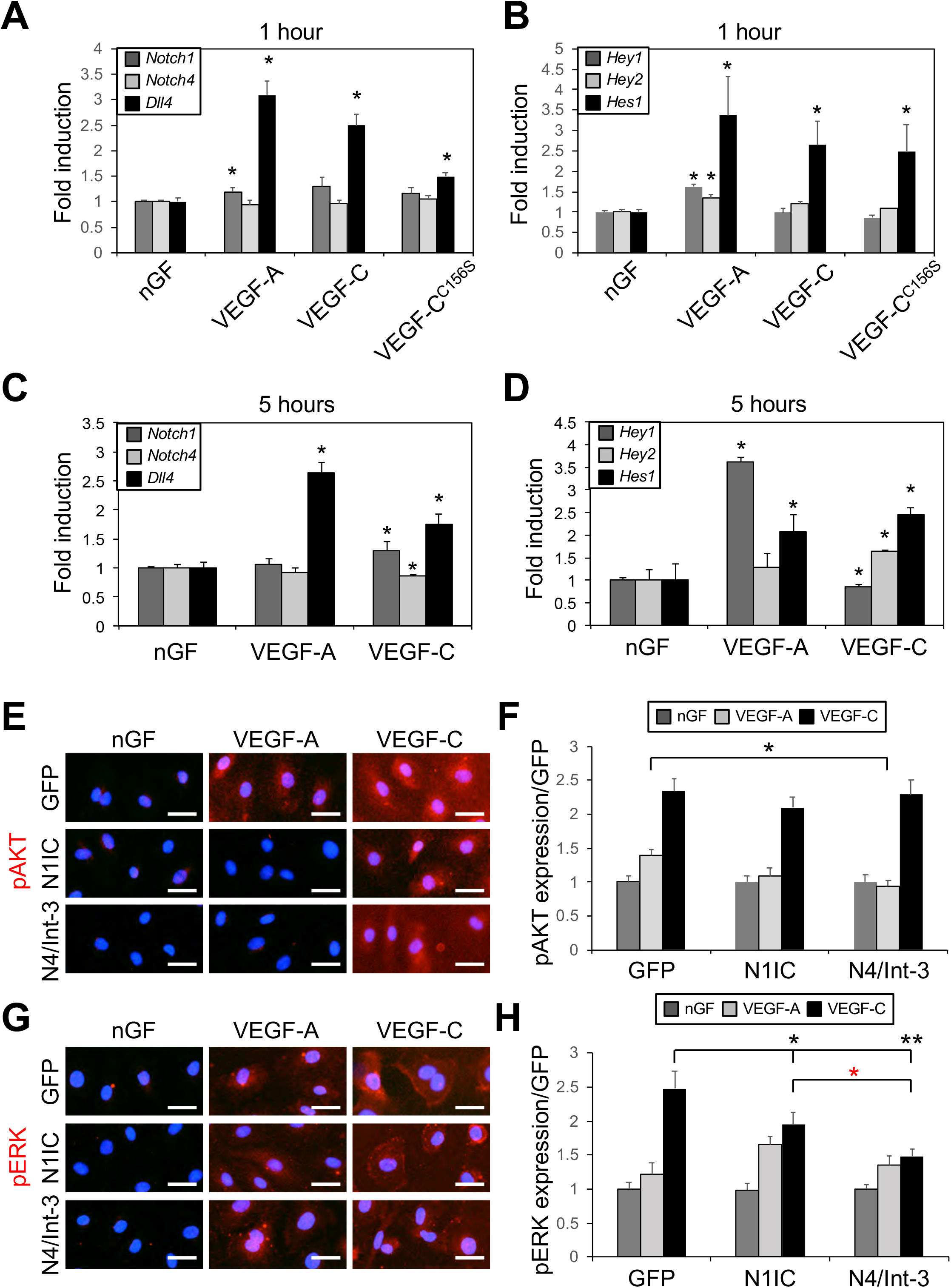
VEGF and NOTCH signaling modulated each other in LECs. **a)** *Notch1, Notch4*, and *Dll4* expression, and **b**) *Hey1, Hey2*, and *Hes1* expression determined by qRT-PCR of HdLECs treated for 1 hour with no growth factor (nGF), VEGF-A, VEGF-C or VEGF-C^C156S^. Experiment done in triplicate. Data presented as mean fold induction relative to HdLECs treated with no growth factor ± s.e.m. *p< 0.05 HdLEC treated with VEGFs relative to HdLEC treated with no growth factor. **c)** *Notch1, Notch4*, and *Dll4* expression, and **d**) *Hey1, Hey2*, and *Hes1* expression determined by qRT-PCR of HdLECs treated for 5 hours with nGF, VEGF-A, or VEGF-C. Data presented as fold induction relative to GFP-expressing HdLEC ± s.d. * p< 0.05 VEGF treated HdLEC relative to HdLEC treated with no growth factor. **e-h**) GFP-, N1IC- and N4/Int-3-HdLECS were treated with VEGF-A or VEGF-C for 20 minutes and then stained for either **e)** phospho-AKT (pAKT), or **g)** phospho-ERK (pERK). Scale bars, 25 µm. Quantification of mean **f)** pAKT and **h)** pERK expression normalized by area. Mean data presented for two independent transductions and experiment performed in duplicate. Data presented as fold expression relative to GFP-expressing HdLECs treated with no growth factor ± s.e.m. **f)** pAKT: two-way ANOVA: p <0.0001, Dunnett’s multiple comparison test *p< 0.002 VEGF-A-treated N4/Int-3 vs GFP HdLEC. **h)** pERK: two-way ANOVA: p <0.001, Dunnett’s multiple comparison test *p< 0.03, **p <0.0001 VEGF-C-treated N1IC and N4/Int-3 vs GFP HdLEC. 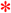 VEGF-C-treated N4/Int-3 vs N1IC HdLEC.

To determine if VEGF-A and VEGF-C differentially regulate Notch signaling in BECs and LECs, HdLECs and HUVEC were serum starved and treated with either VEGF-A or VEGF-C. After 5 hours, VEGF-A only induced *Dll4*, which was associated with an increase in *Hey1* and *Hes1* in HdLECs (Fig. 4c, d). VEGF-C induced *Dll4* in HdLECs (Fig. 4c), which correlated with an increase in *Notch1* transcripts (Fig. 4c), as well as *Hey2* and *Hes1* induction (Fig. 4d). In HUVEC, VEGF-A induced *Dll4* and *Notch1* expression, whereas VEGF-C modestly decreased *Dll4, Notch1* and *Notch4* transcripts (Fig. S4a). Thus, VEGF-A and VEGF-C dynamically and temporally induced Notch activity and specific Notch effectors via induction of *Dll4* in cultured LECs.

To determine if Notch1 or Notch4 activation altered signaling downstream of the VEGFs, HdLECs expressing either N1IC, N4/Int-3 or GFP were serum starved overnight and then stimulated with VEGF-A or VEGF-C. After 20 minutes, HdLECs were assessed for AKT and ERK activation by immunofluorescent staining for phospho-AKT and phospho-ERK (Fig 4e-h.). As compared to the GFP controls, activation of AKT was reduced in N4/Int-3 HdLECs treated with VEGF-A. Activation of AKT by VEGF-C was unaffected in HdLECs expressing either N1IC or N4/Int-3. Levels of ERK activity were unaffected by constitutive activation of Notch1 or Notch4 in all VEGF-A treated HdLECs (Fig. 4e, f). Whereas, ERK activation was reduced in N1IC and N4/Int-3 HdLECs treated with VEGF-C relative to controls (Fig. 4g, h). Previous studies have shown that Notch signaling alters the expression of the VEGF-A and VEGF-C receptors, VEGFR2 and VEGFR3 [41,18,12,9]. The reduction in AKT and ERK activation downstream of VEGFs may be secondary to Notch signaling effects on VEGFR expression. Therefore, the expression of *Vegfr2* and *Vegfr3* was determined in HdLECs with N1IC and N4/Int-3 by qRT-PCR. Both N1IC and N4/Int-3 downregulated *Vegfr2*, while they induced *Vegfr3* (Fig. S4b). Taken together the data suggests that the decrease in AKT activation by VEGF-A and ERK activation by VEGF-C may be due to reduced VEGFR2 levels, and not VEGFR3.

### Embryonic dermal lymphangiogenic defects in Notch4 mutant mice

Prior reports have shown that loss of LEC *Notch1* in embryos leads to increased LEC proliferation and tip cells [14]. To determine the role of Notch4 in embryonic lymphangiogenesis, we evaluated *Notch4*^*-/-*^ mice and compared their lymphatic phenotype to that of *Notch4*^*+/-*^ and wild-type littermates. To confirm that NOTCH4 protein is absent in the *Notch4*^*-/-*^ embryos, Western blots using lysates collected from E14.5 embryos and staining of P4 dermal tissue with an antibody against the intracellular domain of NOTCH4 were performed. As compared to wild-type littermates, NOTCH4 expression was absent in *Notch4*^*-/-*^ tissues (Fig. S5a-c). NOTCH1 expression determined by immunostaining of P4 dermis was unaffected (Fig. S5d).

Analysis of E14.5 *Notch4*^*-/-*^ dermal wholemounts revealed that the distance between the two migrating lymphatic fronts was decreased relative to wild-type and *Notch4*^*+/-*^ littermates (Fig. 5a, b). This correlated with an increase in the number of lymphatic fronts migrating towards the midline in the *Notch4*^*-/-*^ dermis (Fig. 5c). Further analysis of the lymphangiogenic sprouts at the migration front revealed the length from the front to the first branch-point did not differ between mutants and controls (Fig. 5d). We next evaluated the lymphatic vessel caliber at the lymphangiogenic front and in the maturing plexus. The caliber of the vessel adjacent to the first branch-point at the front did not differ between mutant and control mice (Fig. 5e, f). However, a significant reduction of vessel caliber was observed in the maturing lymphatic plexus of *Notch4*^*-/-*^. Although the lymphatic vessel diameter was reduced in the maturing *Notch4*^*-/-*^ plexus, branching was similar between mutants and controls (Fig. S6a). To determine if the reduce dermal lymphatic vessel caliber was due to a change in LEC proliferation, wild-type and *Notch4*^*-/-*^ E14.5 dermal skin were stained for the proliferation marker, KI67, and LYVE1. Dermal LEC proliferation was similar between *Notch4*^*-/-*^ and control mice (Fig. S6b, c). As NOTCH4 is also expressed by the blood vasculature [24,20], we evaluated the underlying dermal blood vascular network in E14.5 *Notch4*^*-/-*^ and *Notch4*^*+/-*^ embryos. Consistent with prior studies, the density and branching of the blood vasculature were unaffected in the *Notch4* nulls (Fig. S7) [42,24].

**Fig. 5.**
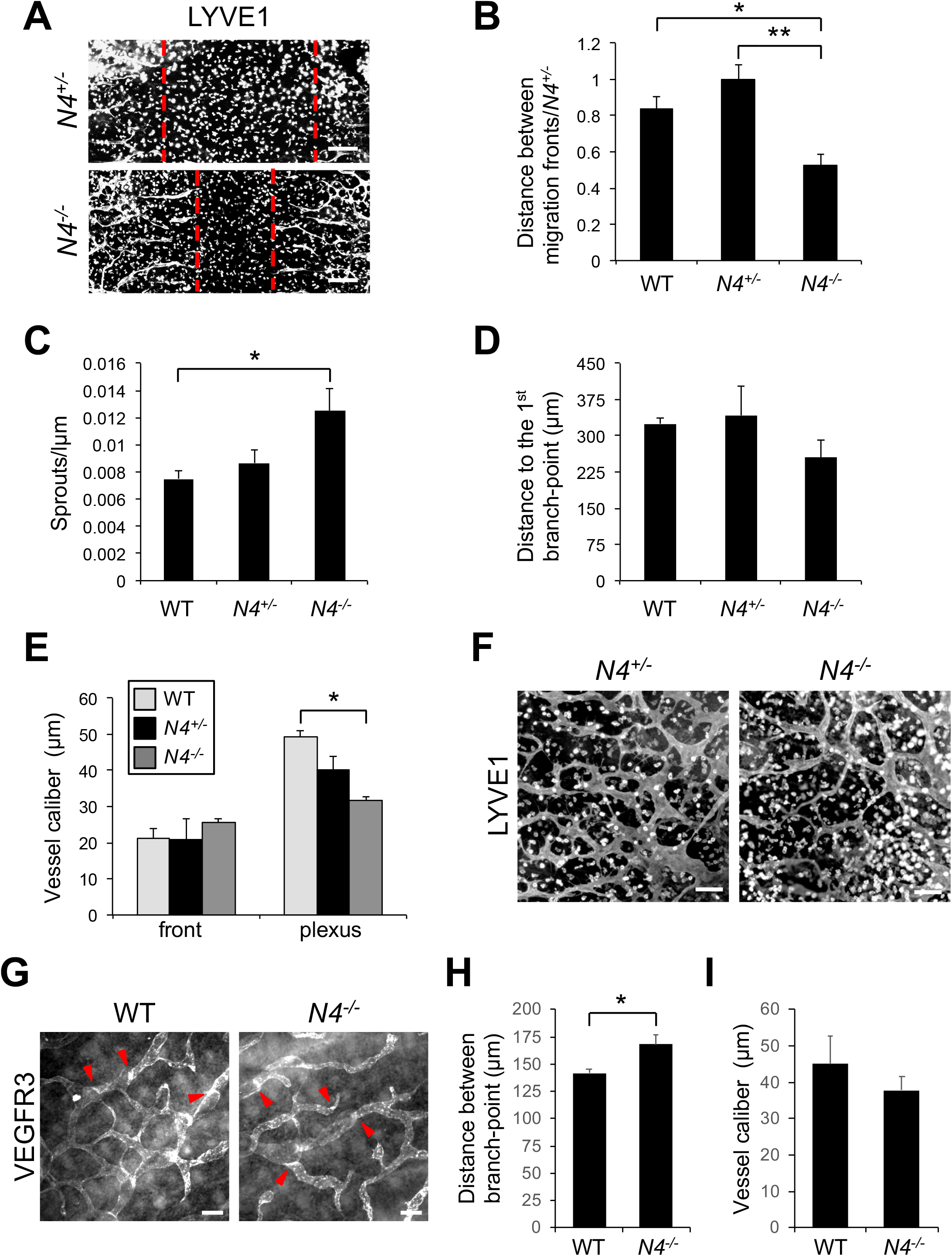
Loss of *Notch4* altered embryonic dermal lymphangiogenesis. **a-f)** Dermal lymphatic phenotype was determined for E14.5 *Notch4*^*-/-*^ (*N4*^*-/-*^), *Notch4*^*+/-*^ (*N4*^*+/-*^) and wild-type (WT) wholemounts stained for LYVE1. **a**) Representative image of *N4*^*-/-*^ and *N4*^*+/-*^ dermal wholemounts. Red dotted line marks leading edge of lymphangiogenic front. Scale bars, 100µm **b**) Quantification of the distance between migration fronts, normalized to the *N4*^*+/-*^ controls. Data presented ± s.e.m. One-way ANOVA: p= 0.014, T-Test: *p< 3 x10^−4^, **p< 0.001. wt (n=5), *N4*^*+/-*^ (n=13), *N4*^*-/-*^ (n=4) **c**) Quantification of the number of lymphangiogenic sprouts normalized by length of the front. Data presented ± s.e.m. One-way ANOVA: p= 0.023, T-Test: *p< 0.02. WT (n=6), *N4*^*+/-*^ (n=11), *N4*^*-/-*^ (n=6) **d**) Quantification of the sprout length from the migration front to first branch-point. Data presented ± s.e.m. WT (n=7), *N4*^*+/-*^ (n=8), *N4*^*-/-*^ (n=6) **e**) Quantification of the average vessel caliber at the lymphangiogenic front and in the maturing plexus. Data presented ± s.e.m. One-way ANOVA: p= 0.019, T-Test: *p< 0.03, front analysis-WT (n=7), *N4*^*+/-*^ (n=13), *N4*^*-/-*^ (n=7), plexus analysis - WT (n=6), *N4*^*+/-*^ (n=9), *N4*^*-/-*^ (n=8) **f**) Representative image of the E14.5 maturing lymphatic plexus in *N4*^*-/-*^ and *N4*^*+/-*^ littermates. Scale bars, 100µm **g-i)** Dermal lymphatic phenotype determined for E16.5 *Notch4*^*-/-*^ (*N4*^*-/-*^) and wild-type (WT) wholemounts stained for VEGFR3. **g)** Representative image of the E16.5 dermal lymphatic plexus in *N4*^*-/-*^ and WT littermates. Red arrowheads mark lymphatic valves. Scale bars, 100µm **h)** Quantification of mean distance between branch-points. Data presented ± s.e.m. T-Test: *p< 0.03 wt (n=4), *N4*^*-/-*^ (n=6) **i)** Quantification of the average vessel caliber of the dermal lymphatic plexus at E16.5. Data presented ± s.e.m. wt (n=6), *N4*^*-/-*^ (n=8)

We next evaluated the dorsal dermal lymphatic phenotype at E16.5 in wild-type and *Notch4*^*-/-*^ embryos. The lymphatic fronts had reached the midline and merged in both wild-type and *Notch4*^*-/-*^ embryos. Analysis of the lymphatic plexus revealed that it was disorganized with an increase in the distance between branch-points, while there was no difference in the mean vessel caliber in *Notch4*^*-/-*^ dermis relative to controls (Fig. 6g-i). Lymphatic valves were observed in both mutants and controls. A recent study has shown that reduced branching in the dermal lymphatic plexus was associated with an increase in blunt-end sprouts due to reduced VEGFR3 signaling, which in turn led to a less branched network [43]. In contrast, increased VEGFR3 signaling was associated with reduced blunt-ended sprouts and a more densely branched network [44]. Since we observed a decreased in the dermal lymphatic branching in the E16.5 *Notch4*^*-/-*^ embryos, we assessed the sprout phenotype at E14.5 during active lymphangiogenesis. The sprouts of control embryos uniformly expressed LYVE1 and were elongated with numerous filopodia consistent with a lymphangiogenic phenotype (Fig. S8). In contrast, LECs in sprouts in the *Notch4*^*-/-*^ lymphatic vasculature were often rounded with reduced and blunted filopodia.

**Fig. 6.**
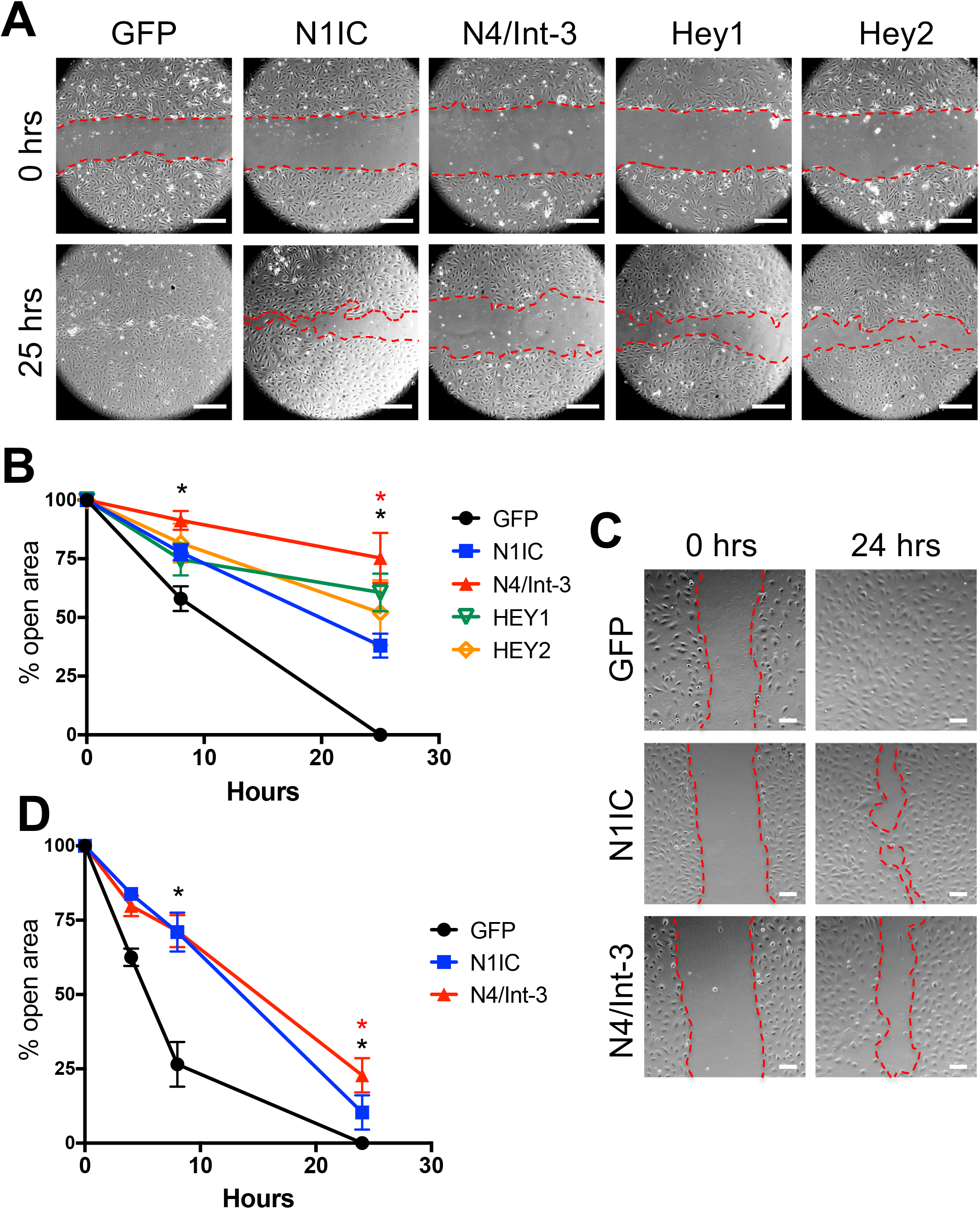
Notch signaling inhibited LEC migration. **a**) Confluent N1IC-, N4/Int-3-, Hey1-, Hey2-, or GFP-expressing HdLECs were scratched and representative images for 0 and 25 hours presented. Scale bars, 2.5µm. **b**) Quantification of percent open wound v area at 0, 8 and 25 hours. Data presented ± s.e.m. two-way-ANOVA: p <0.0012, T-Test: *p <0.002 N1IC and N4/Int-3 vs GFP at 8 and 25 hours, *p< 0.002 N4/Int-3 vs N1IC at 25 hours **c**) Confluent N1IC, N4/Int-3, or GFP expressing HdLECs were treated with mitomycin C and scratched and representative images for 0 and 24 hours presented. Scale bars, 100µm. **d**) Quantification of percent open wound area at 0, 4, 8 and 24 hours. Data presented ± s.e.m. two-way ANOVA: p <0.0001, Dunnett’s multiple comparison test *p< 0.0001 N1IC and N4/Int-3 vs GFP. T-Test: *p< 0.0001 N4/Int-3 vs N1IC.

Thus, *Notch4* mutant mice had a distinct dermal lymphatic phenotype from that observed in mice with LEC *Notch1* deletion [14]. Rather than increased vessel diameter due to increased proliferation and branching due to increased sprouting lymphangiogenesis, the embryonic dermal lymphatics in *Notch4* nulls had increased front closure, an early decrease in the lymphatic vessel caliber, and reduced branching without a change in LEC proliferation.

### NOTCH4 activation preferentially inhibited LEC migration

To determine the effects of Notch1 and Notch4 signal activation on LEC migration, a monolayer-wounding assay was performed using HdLECs expressing either N1IC or N4/Int-3. Relative to control GFP-expressing HdLEC, both N1IC and N4/Int-3 expression inhibited LEC migration (Fig. 6). We next evaluated the effect of overexpressing the downstream Notch effectors HEY1 and HEY2 on HdLEC migration. Ectopic expression of HEY1 and HEY2 suppressed migration relative to control HdLECs. Further analysis revealed that N4/Int-3 was a significantly stronger inhibitor of LEC migration than N1IC at 25 hours. To insure the difference in migration was not due to changes in LEC proliferation, N1IC, N4/Int-3 and GFP expressing HdLECs were treated with mitomycin C and LEC migration determined. Similar to the initial migration assay, both N1IC and N4/Int-3 suppressed HdLEC migration with N4/Int-3 suppressing migration significantly more than N1IC at 24 hours (Fig. 6c,d).

### Inhibition of lymphatic endothelial canonical Notch signaling increased dermal lymphatic vessel density

Notch4 has been shown to signal via RBPjκ-dependent (canonical) and RBPjκ-independent (non-canonical) downstream pathways [45-47]. To determine the effects of LEC specific loss of canonical Notch signaling on dermal lymphangiogenesis, we used the inducible *Prox1CreER*^*T2*^ driver to induce expression of a *DNMAML* transgene [26]. *DNMAML* encodes a dominant negative form of Mammalian Mastermind-like 1 (MAML1) that binds the NOTCH/RBPjκ complex to form an inactive complex and blocks the recruitment of transcriptional co-activators. *Prox1CreER*^*T2*^ mice were crossed with *DNMAML*^*fl/fl*^ mice to generate *Prox1CreER*^*T2*^;*DNMAML*^*fl/+*^ embryos (*DNMAML*^*LEC*^) and *DNMAML*^*fl/+*^ control littermates. To circumvent effects on early lymphatic specification caused by loss of Notch signaling in LECs [18], tamoxifen was administered to pregnant females at E12.5, just as sprouting lymphangiogenesis begins, and the dermal lymphatic phenotype analyzed at E14.5. Unlike the *Notch4* nulls, the closure of the migration fronts was the same between *DNMAML*^*LEC*^ and control (Fig. 7a,b). The number of sprouts along the migrating front and the length of the sprout to the first branch-point were similar between mutants and controls (Fig. 6c, d). In contrast, the lymphatic density was nearly 25% greater in the *DNMAML*^*LEC*^ compared to controls (Fig. 6e). The increase in the *DNMAML*^*LEC*^ dermal lymphatic density correlated with an enlargement of the lymphatic vessel caliber at the lymphangiogenic front and in the maturing plexus (Fig. 7f, g). As compared to controls, *DNMAML*^*LEC*^ dermal lymphatics had an increase in the number Prox1+/LYVE1+ LECs (Fig. 6h), while branching in the mature plexus was unaffected (Fig. 6i). The increase in vascular density was specific to the lymphatics as blood vessel density was unchanged in *DNMAML*^*LEC*^ mutants (Fig. S9). Thus, we found that inhibition of RBPjκ-dependent Notch signaling resulted in increased lymphatic vessel density and caliber associated with an increase in LECs, suggesting canonical Notch signaling suppressed LEC proliferation in the embryonic dermal lymphatics.

**Fig. 7.**
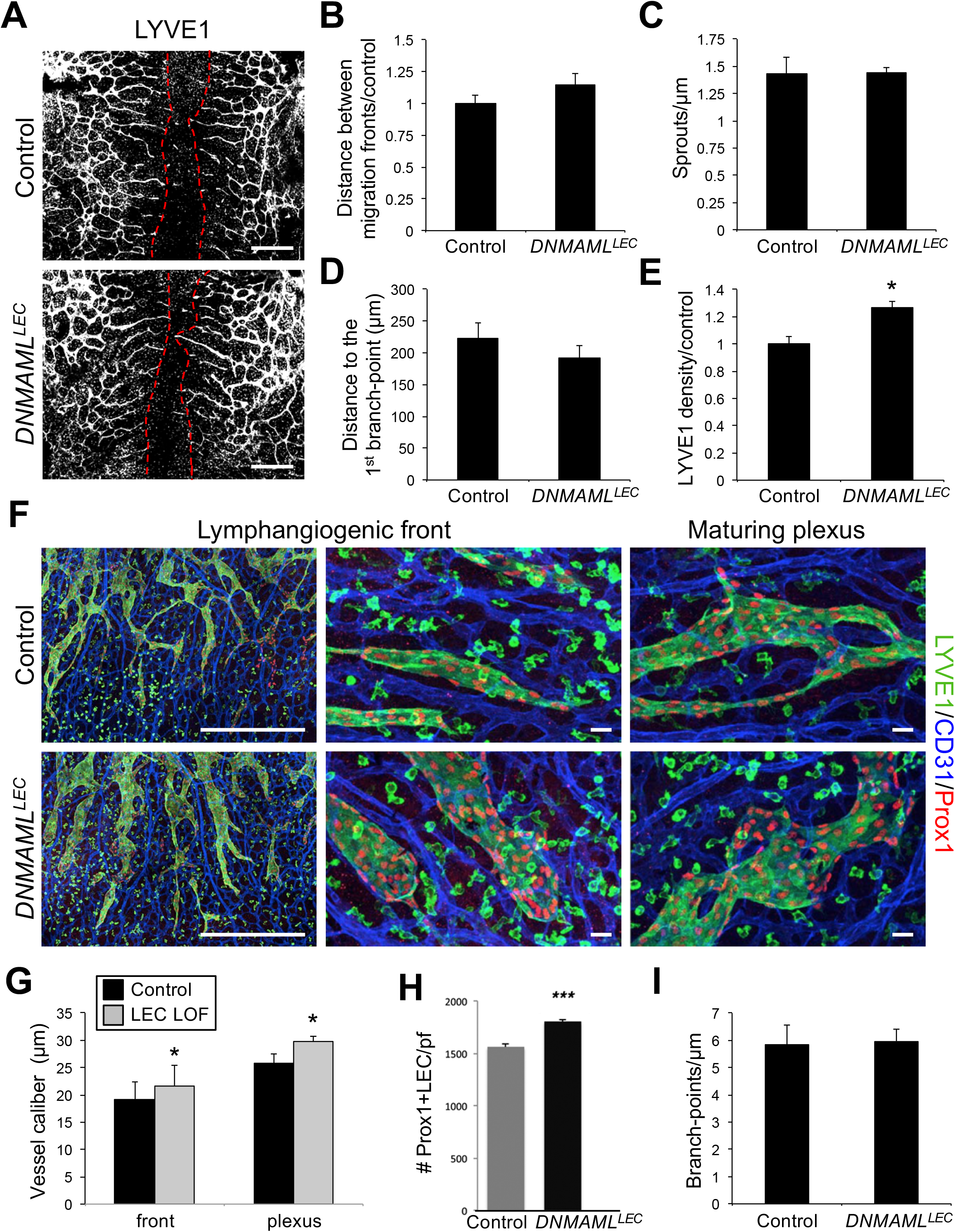
Loss of canonical Notch signaling in LECs increased dermal lymphatic density. *Prox1CreER*^*T2*^ and *DNMAML*^*fl/fl*^ mice were crossed and tamoxifen administered at E12.5 and dorsal dermis analyzed at E14.5. **a**) Representative image of LYVE1 staining of *Prox1CreER*^*T2*^;*DNMAML*^*fl/+*^ (*DNMAML*^*LEC*^) and *DNMAML*^*fl/+*^ (control) dermis. Red dotted line marks leading edge of lymphatic fronts. Scale bars, 1000µm **b**) Quantification of the distance between migration fronts, normalized to the control. Data presented ± s.e.m. Control (n=7), *DNMAML*^*LEC*^ (n=9) **c**) Quantification of the number of lymphangiogenic sprouts normalized by length of the front. Data presented ± s.e.m. Control (n=8), *DNMAML*^*LEC*^ (n=6) **d**) Quantification of the distance from the migration from to first branch-point. Data presented ± s.e.m. Control (n=8), *DNMAML*^*LEC*^ (n=6) **e**) Quantification of average LYVE1+ vessel density normalized by area. Data presented relative to control ± s.e.m. T-Test * p<0.002, control (n=7), *DNMAML*^*LEC*^ (n=9) **f**) LYVE1, CD31 and PROX1 staining of *DNMAML*^*LEC*^ mutant and control dermal wholemounts. Images represent low (left) and high (middle) magnification of the lymphangiogenic front and the maturing plexus (right). Scale bars, 500µm (left), 100µm (middle, right) **g**) Quantification of the average vessel caliber at the lymphangiogenic front and in the maturing plexus. Data presented ± s.e.m. T-Test: *p< 0.04, **p<0.01, control (n=8), *DNMAML*^*LEC*^ (n=6) **h**) Quantification of the number of Prox1+ LECs per field (pf). Data presented as ± s.d. ***p<0.001. control (n=8), *DNMAML*^*LEC*^ (n=6) **i**) Quantification of the average number of branch-points normalized to unit of vessel length. Data presented ± sem. control (n=3), *DNMAML*^*LEC*^ (n=5)

### Notch4^-/-^ and DNMAML^LEC^ embryos display distinct lymphatic phenotypes at E17.5

To assess the functionality and patterning of the dermal lymphatics, lymphangiographies were performed on E17.5 *Notch4*^*-/-*^, *DNMAML*^*LEC*^ and control littermates. Dye was injected within the dermis in the periorbital region and uptake by the lymphatics assessed after 1 minute. The *Notch4*^*-/-*^ lymphatic plexus had reduced branching with tortuous vessels relative to the more uniform lymphatics of controls (Fig. 8a). One of 8 *Notch4*^*-/-*^ embryos analyzed had blood-filled dermal lymphatics at E17.5 (Fig. 8b). In contrast to the *Notch4*^*-/-*^ phenotype, *DNMAML*^*LEC*^ dermal lymphatics were dilated relative to control littermates (Fig. 8c). One out of 5 *DNMAML*^*LEC*^ embryo lymphatics were leaky (Fig. 8d), which was not observed in controls or *Notch4*^*-/-*^ lymphangiographies. These data demonstrate that the *Notch4* null and mice with a loss of LEC RBPjκ-dependent Notch signaling have distinct phenotypes at E17.5, as well as at E14.5 (Fig. 5, 7).

**Fig. 8.**
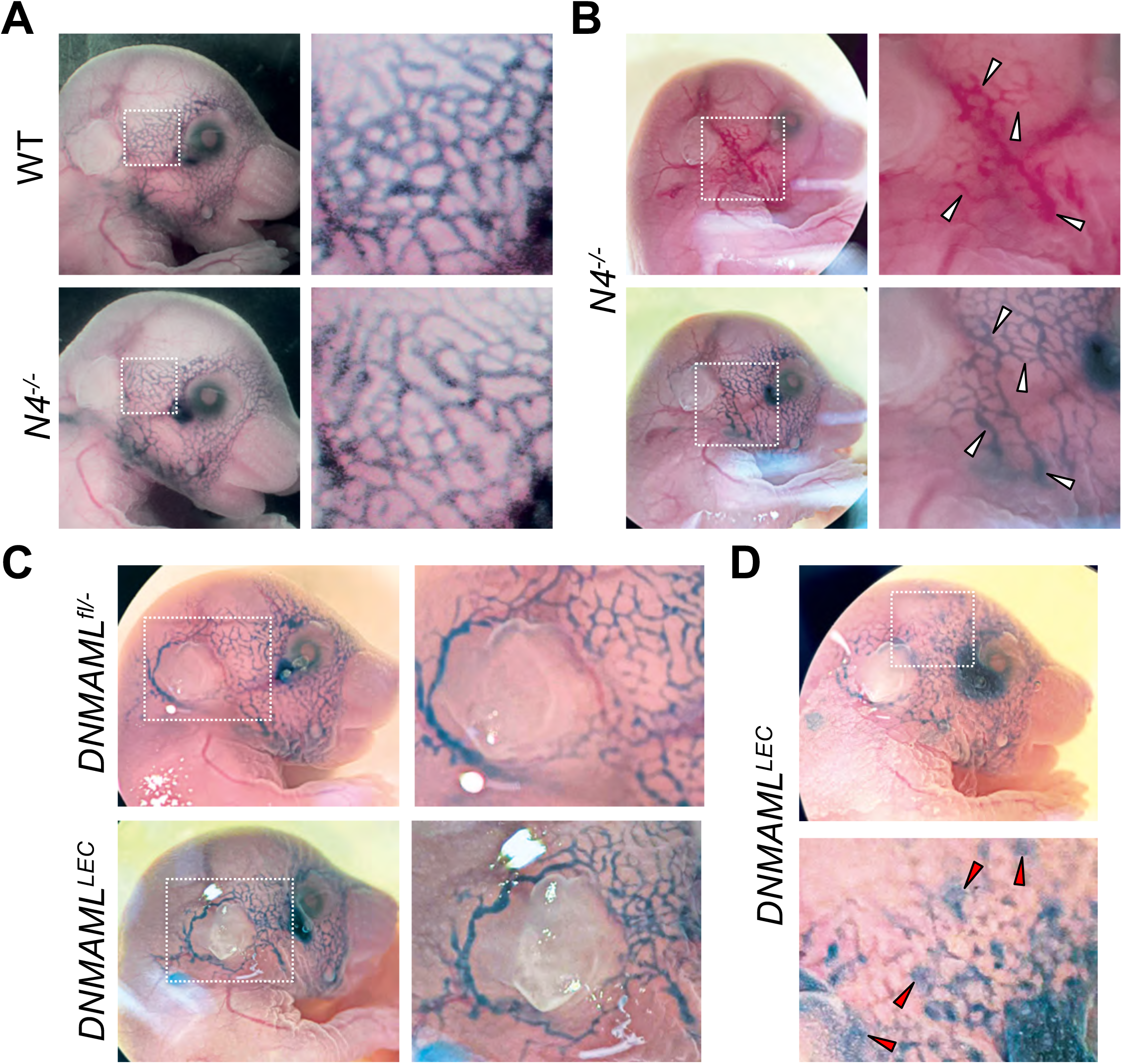
Loss of *Notch4* and deletion of LEC canonical Notch signaling resulted in distinct lymphatic phenotypes at E17.5. **a, b)** Lymphangiography of E17.5 wild-type (n=6) and *Notch4*^*-/-*^ (n=8) embryos. Representative images of wild-type and *Notch4*^*-/-*^ embryos. Boxed area enlarged to the right. **b)** *Notch4*^*-/-*^ embryos with blood-filled dermal lymphatics (white arrowheads). Boxed area enlarged to the right. **c, d)** *Prox1CreER*^*T2*^ and *DNMAML*^*fl/fl*^ mice were crossed and tamoxifen administered at E12.5 and lymphangiography performed at E17.5. Control (n=12), *DNMAML*^*LEC*^ (n=5) **c)** Representative images of *DNMAML*^*fl/-*^ control and *DNMAML*^*LEC*^ embryos. Boxed area enlarged to the right. d) *DNMAML*^*LEC*^ embryo with leaking dermal lymphatic vessels (red arrowheads). Boxed area enlarged to the right.

### Canonical Notch signaling is unaffected in Notch4^-/-^ embryonic dermal lymphangiogenesis

As we observed a difference between the embryonic dermal lymphatic phenotypes of *Notch4*^*-/-*^ and *DNMAML*^*LEC*^ mutants, we evaluated canonical Notch signaling by introducing the *NVR* and *ProxTom* alleles into the *Notch4* null background. Loss of *Notch4* did not change canonical Notch signaling at the lymphangiogenic vascular front, nor the maturing lymphatic plexus (Fig. 9, S6d). This data suggested that Notch4 is not necessary for canonical Notch signaling in the embryonic dermal lymphatics.

**Fig. 9.**
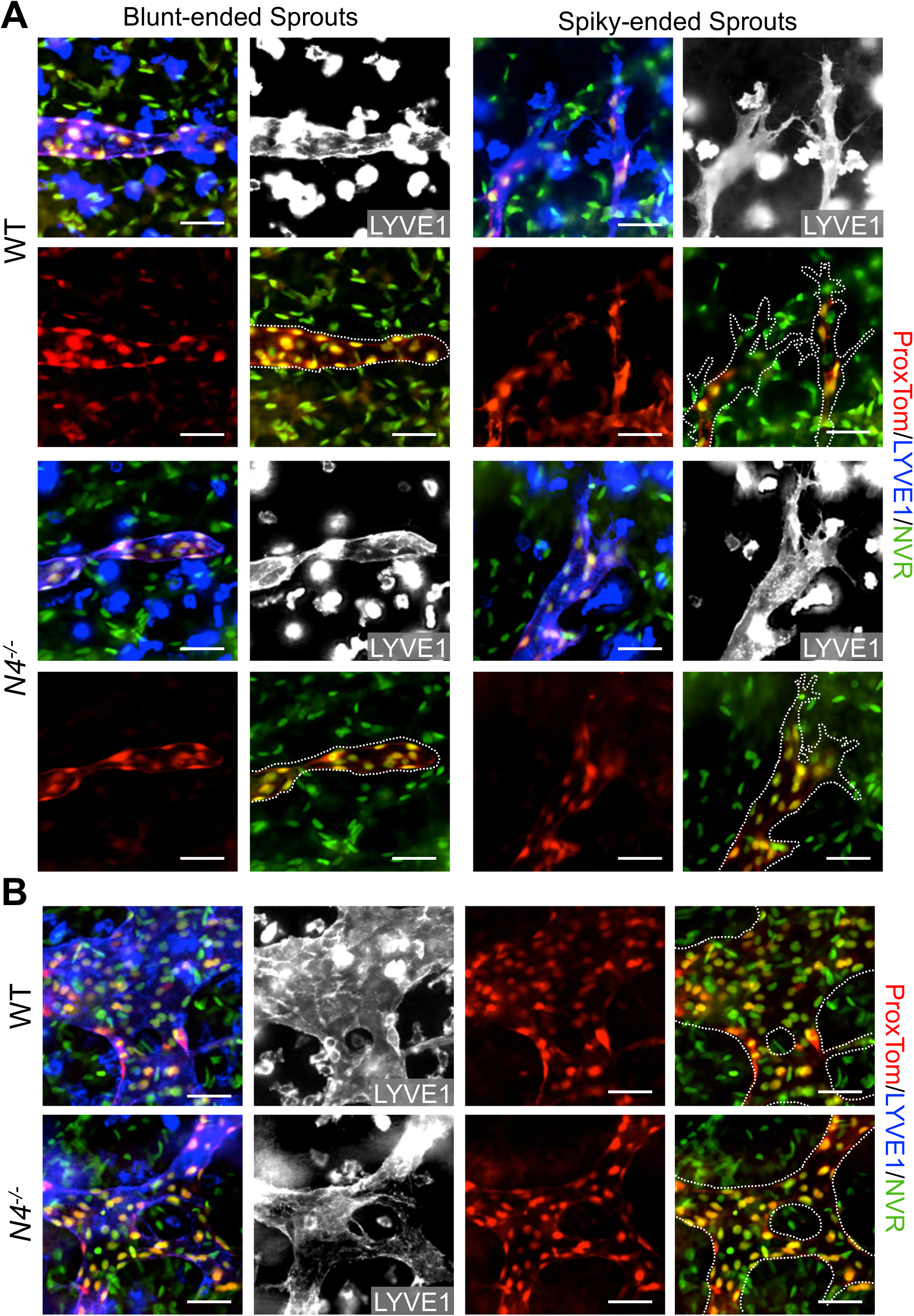
Canonical Notch signaling in LECs is unaltered in the *Notch4* mutant dermal lymphatics. *Notch4*^*+/-*^;*ProxTOM*^*+/-*^;*NVR*^*+/-*^ males were bred with *Notch4*^*+/-*^;*ProxTOM*^*+/-*^;*NVR*^*+/-*^ females, and E14.5 *ProxTOM;NVR* and *Notch4*^*-/-*^;*ProxTOM;NVR* dermal wholemounts stained for LYVE1. **a**) Representative images of blunted-ended and spiky-ended sprouts at the lymphangiogenic front of wild-type and *Notch4*^*-/-*^ dermis. **b**) Representative images of branch-points with Notch signaling in wild-type and *Notch4*^*-/-*^ in the maturing lymphatic vascular plexus. Scale bars, 50µm.

## Discussion

NOTCH1 and NOTCH4 are expressed and Notch signaling active in the embryonic and early postnatal dermal lymphatic vasculature [14,12], suggesting a role for both Notch proteins in embryonic lymphangiogenesis. Loss of *Notch4* was shown to exacerbate the *Notch1* null embryonic blood vascular phenotype, suggesting Notch1 and Notch4 have overlapping functions in the blood endothelium [24]. In contrast to the blood endothelium, we found that loss of *Notch4* led to a distinct embryonic dermal lymphangiogenic phenotype, then that observed in mice with LEC deletion of *Notch1* [14], or inhibition of canonical Notch signaling, presented here. At E14.5, *Notch4* null embryos displayed an increase in the closure of the lymphangiogenic fronts to the midline, reduced vessel caliber in the maturing plexus, and an increase in blunt-ended sprouts, while LEC proliferation was unaffected. By E16.5, the dermal lymphatic plexus in *Notch4*^*-/-*^ has reduced branching and tortuous lymphatic vessels, which may be secondary to the increase in blunt-ended sprouts at E14.5. In cultured LECs, constitutive activation of Notch4 was a stronger inhibitor of migration than Notch1 activation and induced of a subset of lymphangiogenic genes. In contrast, loss of LEC *Notch1* at E10.5 increased embryonic dermal lymphatic density, due to increased LEC proliferation and decreased LEC apoptosis [14]. Similar to the *Notch1* LEC knockout, we demonstrate that LEC expression of DNMAML, which inhibits canonical Notch/RbpJκ signaling, increased the dermal lymphatic vascular density consistent with an increase in LEC proliferation and viability. Distinct functions for Notch1 and Notch4 have been described for endothelial progenitor cells, where Dll4 signaling via Notch4 specifically induced EphrinB2 and increased proliferation and migration of cultured cells [48]. More recently, it was proposed that endothelial Dll4/Notch1 signaling induces Hey2 to suppress proliferation and tip cell formation, while Jag1 activates Notch4 to induce Hey1 and promote vessel maturation while having no effect on vascular density [49]. Taken together, we propose that Notch1 and Notch4 signal dynamically to regulate lymphangiogenesis and control migration and branching, versus proliferation and cell viability by distinct mechanisms.

Our studies suggest that Dll4 signaling via Notch1 and Notch4 have overlapping and unique transcriptional targets in HdLECs. Constitutive Notch1 and Notch4 activation in LECs both induced the expression of Notch effectors (*Hes1, Hes4, Hey1, Hey2, HeyL*) and lymphangiogenic genes, such as *EphrinB2*, and *Cx37* [50], as well as downregulated essential genes in lymphangiogenesis, *Podoplanin*, and *Prox1*. In contrast, *Hes5* expression was induced by N1IC, and not N4/Int-3. DLL4/Notch induced genes involved in chemokine signaling were also differentially regulated by Notch1 and Notch4. Expression of *Cxcr4* was induced by Notch1 activation, but suppressed by Notch4 signaling. In LECs, CXCR4 signaling promoted wound-induced and VEGF-C driven lymphangiogenesis in vivo, while in vitro it induced chemokine-driven LEC migration [31,38]. Thus, it is possible that loss of *Notch4* led to an increase in CXCR4 expression, which in turn increased the LEC migration towards the midline. N4/Int-3 was also a significantly stronger inducer of *Ccl2* and *Ackr3*, than N1IC. LEC-derived CCL2 has been shown to promote the recruitment of monocytes and macrophages to sites of lymphangiogenesis, where they deliver VEGF-A and VEGF-C [33,36]. In murine lymphatic development, ACKR3 functions to suppress LEC growth by scavenging adrenomedullin (ADM) [35]. Interestingly, we found that Dll4/Notch signaling also suppressed *Adm* expression, suggesting Dll4/Notch4 signaling suppresses ADM signaling to regulate lymphatic development. While our gene expression studies begin to elucidate some of the mechanisms by which Dll4/Notch1 and Dll/Notch4 signaling regulates lymphangiogenesis, further studies are necessary to understand the complexity of Notch1 and Notch4 signaling in LECs.

Prior studies have shown that VEGF-C induces DLL4 in LECs leading to Notch activation [10]. We expanded these studies to understand the role of time and specific VEGFRs in this process. By using VEGF-C^C156S^ which specifically binds VEGFR3 and VEGF-A which binds VEGFR2, we found that VEGFR2 signaling was a stronger and faster inducer of Dll4/Notch signaling than VEGFR3 signaling. We also discovered that the induction of the *Hes* and *Hey* genes was time- and VEGF-dependent in HdLECs. VEGF-A induced *Hes1* at 1 hour which persisted until 5 hours, while significant *Hey1* upregulation was not observed until 5 hours. In LECs, VEGF-A was a stronger inducer of *Hey1*, while VEGF-C induced *Hey2*. This differential response of *Hes* and *Hey* genes to VEGF-A and VEGF-C was specific to LECs, as VEGF-C had no effect on *Dll4*, or Notch gene expression in HUVEC. Our studies also suggest that additional *Hes* and *Hey* gene family members than those studied in the blood vasculature, *Hes4* and *HeyL*, have a role in transmitting Notch signaling in LECs.

Our studies also revealed that Notch1 and Notch4 differentially altered signaling downstream of VEGF-A/VEGFR and VEGF-C/VEGFR. Constitutive activation of Notch4 blocked AKT activation by VEGF-A, and ERK activation by VEGF-C. whereas Notch1 activity only modestly suppressed VEGF-C activation of ERK signaling. Together these data suggest that Notch4 has a role in modulating VEGFR2 and VEGFR3 signaling down the PI3K/AKT and RAS/MAPK pathway.

The dermal lymphatic phenotypes were distinct between *Notch4*^*-/-*^ and *DNMAML*^*LEC*^, suggesting Notch4 signals at least in part via a non-canonical pathway. Notch4 has been shown to signal via canonical (RBPjκ-dependent) and non-canonical (RBPjκ-independent) Notch pathways in multiple cells types [45-47]. In endothelial cells, Notch4 activation blocked LPS induced apoptosis via RBPjκ-independent upregulation of Bcl2 [45]. In mice, NOTCH4 activation in the ductal epithelium required RBPjκ for physiological alveolar development, but not for breast cancer development, suggesting Notch4 functions via both canonical and non-canonical pathway in the breast endothelium [46,47]. We observed that canonical Notch signaling was unchanged in the embryonic dermal LECs in *Notch4* nulls suggesting the Notch4 dermal lymphatic phenotype did not occur via a RBPjκ-dependent mechanism. However, it possible that the variable phenotypes are due to differences in the penetrance of global *Notch4* loss versus a tamoxifen-induced cell mosaic expression of *DNMAML* in LECs.

An increase in the closure of the two lymphangiogenic fronts was observed in *Notch4* mutants that correlated with reduced vessel caliber in the absence of a change in LEC proliferation. This phenotype is consistent with an increase in LEC migration towards the midline. In HdLECs, ectopic Notch4 activation inhibited LEC migration significantly more than Notch1 activation. This inhibition of LEC migration by Notch4 may occur via non-canonical Notch signaling, as expression of DNMAML, an inhibitor of canonical Notch signaling did not affect the closure of the lymphangiogenic fronts. Notch4 may suppress LEC migration via its interactions with Wnt/β-catenin signaling. Non-canonical Notch4 signaling has been shown to antagonize Wnt/β-catenin signaling in stem and progenitor cells [51,29]. In LECs, loss of β-catenin signaling reduced LEC migration towards the midline and increased dermal lymphatic vessel caliber [51,29], phenotypes opposite to that observed in *Notch4*^*-/-*^ embryos, suggesting that Notch4 via a non-canonical signaling suppresses LEC migration.

Western analysis of embryo lysates and immunostaining of tissue sections using an antibody against the cytoplasmic domain of NOTCH4 demonstrated a loss of NOTCH4 expression in the *Notch4* nulls. It has been suggested that this *Notch4* null line expresses a truncated extracellular NOTCH4 peptide that suppresses Notch1 signaling by functioning as a ligand trap [52]. However, a loss of canonical Notch signaling was not observed in the lymphatics of *Notch4* mutant mice, which would be predicted if Notch1 signaling was inhibited in the model. Moreover, the *Notch4* mutant dermal lymphatic phenotype is distinct from that observed in mice with *Notch1* deleted in the LECs [14], as well as the *DNMAML*^*LEC*^ mice. The dermal lymphatic phenotype however may be due to loss of *Notch4* in non-LECs, such as macrophages, and a conditional *Notch4* allele needs to be developed to better understand the cell type specific requirement for NOTCH4 in lymphatic development.

Together with published data, our studies suggest that Notch1 and Notch4 function distinctly embryonic dermal lymphangiogenesis via a RBPjκ-dependent and –independent pathways. We propose that Dll4/Notch1 signaling via a canonical pathway suppresses LEC proliferation, while Notch4 signaling suppresses LEC migration and branching, possibly via a RBPjκ-dependent mechanism. Further studies into the mechanistic interaction between Notch1 and Notch4 in LECs and lymphatic development and homeostasis is necessary, as a number of therapeutics that are pan-Notch inhibitors or target specific receptors or ligands are currently in clinical trials or the research pipeline for use in the clinic.

## Acknowledgements

The authors thank Valeriya Borisenko and Marina Vorontchikhina for technical assistance, June Wu for critical reading of the manuscript, and Warren Pear (*DNMAML*^*fl/fl*^), Tom Gridley (*Notch4*^*-/-*^), Guillermo Oliver (*Prox1CreER*^*T2*^). and Hong Young Kwon (*Prox1-tdTomato*) for providing mice.

## Authorship Contributions

AM and MKU share first authorship. AM, MKU, YM, JKK, CJS contributed to the study conception and design. Material preparation, data collection and analysis were performed by AM, MKU, GSD, BS, JMJ, AM, SWY, JDM, CK, MG, GR, CJS. The first draft of the manuscript was written by AM, MKU and CJS, and revised by CJS.

## Compliance with ethical standards

### Ethical approval

Isolation of HUVEC and HdLEC from anonymous discarded specimens and received IRB exempt status by Columbia University IRB (AAAA7338). All procedures performed in studies involving human participants were in accordance with the ethical standards of the institutional and/or national research committee and with the 1964 Helsinki declaration and its later amendments or comparable ethical standards. Mouse studies were approved by Columbia University IACUC (AC-AAAE2653, AC-AAAD0577, AC-AAAP9603, AC-AAAP0452, AC-AABB9551). All procedures performed in studies involving animals were in accordance with the ethical standards of the institution or practice at which the studies were conducted.

