## Supplemental data for "Unique functions for Notch4 in murine embryonic lymphangiogenesis"

**Article Title:** *Unique functions for Notch4 in murine embryonic lymphangiogenesis*

**Journal Name:** *Angiogenesis*

**Corresponding Author:**

Carrie J. Shawber

Supplemental Tables 1-3

**Supplemental Table 1. Primers for quantitative RT-PCR using gene specific standards**

| Gene | Upper Primer | Lower Primer |
|---|---|---|
| *β-actin* | 5' CGAGGCCCAGAGCAAGAGAG 3' | 5' CTCGTAGATGGGCACAGTGTG 3' |
| *Dll4* | 5' CGGGTCATCTGCAGTGACAAC 3' | 5' AGTTGAGATCTTGGTCACAAAACAG 3' |
| *Hes1* | 5' CCCAACGCAGTGTCACCTTC 3' | 5' TACAAAGGCGCAATCCAATATG 3' |
| *Hey1* | 5' ACGAGAATGGAAACTTGAGTT C 3' | 5' AACTCCGATAGTCCATAGCAAG 3' |
| *Hey2* | 5' ATGAGCATAGGATTCCGAGAGTG 3' | 5' GGCAGGAGGCACTTCTGAAG 3' |
| *Notch1* | 5' CTCACCTGGTGCAGACCCAG 3' | 5' GCACCTGTAGCTGGTGGCTG 3' |
| *Notch4* | 5' GGTGACACCCCTGATGTCAG 3' | 5' AGCCTGGCAGCCAGCATC 3' |
| *Lyve1* | 5′ TAGCTTTGAAACTTGCAGCTATG 3' | 5′ TCAACAAATGGTTCAGTTTCTGTAG 3' |
| *Podoplanin* | 5′ CCCAGGAGAGCAACAACTCAAC 3' | 5′ CTCGATGCGAATGCCTGTTAC 3' |
| *Prox1* | 5′ ACGTAAAGTTCAACAGATGCATTAC 3' | 5′ CCAGCTTGCAGATGACCTTG 3' |
| *Vegfr2* | 5′ GGACTGGCTTTGGCCCAAT 3′ | 5′ CTTGCTGTCCCAGGAAATTCTG 3′ |
| *Vegfr3* | 5′ GAGACCTGGCTGCTCGGAAC 3' | 5′ TCAGCATGATGCGGCGTATG 3' |

**Supplemental Table 2. Quantitative RT-PCR primers for mRNA seq validation**

| Gene | Forward Primer | Reverse Primer |
|---|---|---|
| *Ackr3* | 5’ CCAAGACCACAGGCTATGACA ‘3 | 5’ TGGTTGTGCTGCACGAGACTGA ‘3 |
| *β-actin* | 5’ CGAGGCCCAGAGCAAGAGAG ‘3 | 5’ CTCGTAGATGGGCACAGTGTG ‘3 |
| *Bmp2* | 5’ TTCGGCCTGAAACAGAGACC ‘3 | 5’ CCTGAGTGCCTGCGATACAG ‘3 |
| *Ccl2* | 5’ AGAATCACCAGCAGCAAGTGTCC ‘3 | 5’ TCCTGAACCCACTTCTGCTTGG ‘3 |
| *Cxcr4* | 5’ ACGCCACCAACAGTCAGAG ‘3 | 5’ AGTCGGGAATAGTCAGCAGGA ‘3 |
| *Efnb2* | 5’ TATGCAGAACTGCGATTTCCAA ‘3 | 5’ TGGGTATAGTACCAGTCCTTGTC ‘3 |
| *Inhbb* | 5’ CGGGTCCGCCTATACTTCTTC ‘3 | 5’ CGTAGGGCAGGAGTTTCAGG ‘3 |
| *Hes1* | 5’ CCTGTCATCCCCGTCTACAC ‘3 | 5’ CACATGGAGTCCGCCGTAA ‘3 |
| *Hes4* | 5’ GAGCGCGTATTAACGAGAGCCT ‘3 | 5’ CTCACGGTCATCTCCAGGATGT ‘3 |
| *Hes5* | 5’ CTCAGCCCCAAAGAGAAAAA ‘3 | 5’ GACAGCCATCTCCAGGATGT ‘3 |
| *Hey1* | 5’ ATCTGCTAAGCTAGAAAAAGCCG ‘3 | 5’ GTGCGCGTCAAAGTAACCT ‘3 |
| *Hey2* | 5’ GCCCGCCCTTGTCAGTATC ‘3 | 5’ CCAGGGTCGGTAAGGTTTATTG ‘3 |
| *HeyL* | 5’ GGCTGCTTACGTGGCTGTT ‘3 | 5’ GACCCAGGAGTGGTAGAGCAT ‘3 |
| *Sema3g* | 5’ CAGAGGATGGGACCTACGATG ‘3 | 5’ GTTGGCACCTTAAACACCTGG ‘3 |
| *Tgfrb2* | 5’ GTAGCTCTGATGAGTGCAATGAC ‘3 | 5’ CAGATATGGCAACTCCCAGTG ‘3 |
| *Unk5b* | 5’ CTGGGACCTTATGCCTTCAA ‘3 | 5’ CGCTTTGGTGGCAAAGTAAT ‘3 |

**Supplemental Table 3. Antibodies**

| Antigen | Supplier | Catologue # | Use |
|---|---|---|---|
| β-ACTIN | Abclonal | AC038 | Westerns |
| CD31 | Pharmingen | 553370 | wholemount IHC |
| DLL4 | R&D Systems | AF1389 | wholemount IHC |
| JAG1 | R&D Systems | AF599 | wholemount IHC |
| KI67 | Abcam | ab15580 | wholemount IHC |
| LYVE1 | Abcam | ab14917 | wholemount, section IHC |
| LYVE1 | Ebiosciences | 14-0443 | wholemount IHC |
| PROX1 | Angiobio | 11-002 | wholemount IHC |
| NOTCH1 | R&D Systems | AF1057 | section IHC |
| NOTCH1 | Cell Signaling | 3608S | wholemount IHC |
| NOTCH4 | J. Kitajewski | RB2-2 | section IHC, Westerns |
| NOTCH4 | BioXCell | BE0129 | wholemount IHC |
| phospho-AKT | Cell Signaling | 13038 | Cell IHC |
| phospho-ERK | Cell Signaling | 4370 | Cell IHC |
| VEGFR3 | R&D Systems | AF743 | wholemount IHC |

**Supplemental Figures 1-9**

Fig. S1 Murine dermal lymphangiogenesis and NOTCH1 and NOTCH4 expression. **a**) The lymphangiogenic plexus consists of a maturing plexus (1) and active lymphangiogenic fronts (2) which migrates towards the lateral midline (3, dashed line). LYVE1 and CD31 staining of E14.5 dorsal skin. Scale bar, 100µm. **b, c**) Cross-sections of wild-type dermis stained for LYVE1 and **b**) NOTCH1 or **c**) NOTCH4. White arrowheads mark LYVE1+/NOTCH1+ and LYVE1+/NOTCH4+ LECs. Yellow arrowheads mark NOTCH1+ blood vessels. Blue arrowheads mark LYVE1+/NOTCH4+ macrophages. Scale bars, 50µm.

Fig. S2 Dll4 and Jag1 activated Notch signaling in HdLECs. **a**) Quantification of ectopic *Dll4* and *Jag1* transcript levels in HeLa cells expressing GFP, DLL4, JAG1 or both DLL4 and JAG1. *Dll4* and *Jag1* qRT-PCR of HeLa cell lines used in co-culture. Data presented as relative transcript levels ± s.e.m. **b**) Notch/CSL luciferase reporter assay of ligand-expressing or GFP-expressing HeLa cells (y-axis) co-cultured with HdLEC. Data presented as fold luciferase induction over HdLEC co-cultured with GFP-HeLa controls ± s.e.m. Representative experiments presented and performed in triplicate. * $p < 0.02$, ** $p < 0.001$. ns – not significant.

Fig. S3 Effect of DLL4/Notch signaling on Notch pathway and lymphangiogenic genes in HdLECs. **a**) Heatmap of Notch pathway genes. **b**) Heatmap of lymphangiogenic genes.

Fig. S4 VEGF and Notch signaling in HUVEC and HdLECs. **a**) *Notch1*, *Notch4*, and *Dll4* qRT-PCR of HUVEC treated for 5 hours with either VEGF-A or VEGF-C. Data presented as fold induction over HUVEC in no growth factors ± s.d. * $p < 0.05$. b) *Vegfr2*, *Vegfr3*, *Prox1*, *Podoplanin*, and *Lyve1* qRT-PCR of HdLEC expressing GFP, N1IC, or N4/Int-3. Data presented as fold induction over GFP expressing HdLECs ± s.e.m. Mean for > three experiments presented. T-Test * $p < 0.05$.

Fig. S5 Validation of the *Notch4* null mice. **a**) P4 dorsal dermal tissue cross-sections from $Notch4^{-/-}$ and wild-type littermates were stained for VEGFR3 and NOTCH4. NOTCH4 expression was absent in the lymphatic endothelium (white arrowheads) and epithelial cells of the hair follicle (yellow arrowheads) and dermis in $Notch4^{-/-}$ tissues. **b**) Representative Western blot of E14.5 $Notch4^{-/-}$ and wild-type tissues probed with antibodies against the cytoplasmic domain of NOTCH4 (N4IC) or βACTIN. **c**) Mean N4IC expression normalized by βACTIN for wild-type (n=3) and $Notch4^{-/-}$ (n=4) Westerns determined by densitometry. Data presented ± s.e.m. T-test: * $p < 0.05$. **d**) P4 dorsal dermal tissue cross-sections from $Notch4^{-/-}$ and wild-type littermates were stained for VEGFR3 and NOTCH1. NOTCH1 expression was unchanged in the VEGFR3+ lymphatics (white arrowheads). Scale bars, 50µm.

Fig. S6 Lymphatic branching, proliferation and canonical Notch signaling was unaffected in E14.5 $Notch4^{-/-}$ dermis. **a)** Quantification of the average number of branch-points normalized to unit of vessel length. Data presented ± sem. wt (n=7), $N4^{+/-}$ (n=13), $N4^{-/-}$ (n=7). **b**) LYVE1 and KI67 staining of E14.5 $Notch4^{-/-}$ and wild-type dermis. Scale bars, 100µm. **c**) Quantification of PROX1+/KI67+ LECs normalized to lymphatic vascular area ± s.e.m. wt (n=3), $N4^{-/-}$ (n=4) **d**) Number of PROX1+ and PROX1+/NVR+ LECs was determined for individual sprout at the lymphangiogenic front until the first branch-point for E14.5 *ProxTOM;NVR* and $Notch4^{-/-}$*;ProxTOM;NVR* wholemounts stained for LYVE1, Data presented as percent PROX1+/NVR+ LECs relative to total Prox1+ LECs ± s.e.m. wt (n=6), $N4^{-/-}$ (n=3)

Fig. S7 Dermal blood vasculature was unaffected in $Notch4^{-/-}$ mice. **a**) CD31 staining of E14.5 dorsal dermal wholemounts from $N4^{-/-}$ and control $N4^{+/-}$ littermates. **b**) Quantification of average CD31 intensity normalized by area. Data presented ± s.e.m. $N4^{+/-}$ (n=3), $N4^{-/-}$ (n=4). **c**) Quantification of the average number of branch-points per field of view. Data presented ± sem. $N4^{+/-}$ (n=3), $N4^{-/-}$ (n=5).

Fig. S8 Increase in blunt-ended lymphatic sprouts in E14.5 $Notch4^{-/-}$ dermis. **a)** High magnification images of LYVE1+ sprouts at the lymphangiogenic front. White arrowheads mark filopodia extending from a spiky-ended sprouts. Red arrowheads mark blunt-ended sprouts, Scale bars, 20µm. **b)** Quantification of the sprout morphology at the lymphangiogenic front. Blunt-ended sprouts are rounded with reduced filopodia. Spiky-ended sprouts have multiple filopodia. Data presented as percent blunt or spiky phenotype relative to total sprouts ± s.e.m. One-way ANOVA: p= 0.004, T-Test: *$p < 0.003$, wt (n=5), $N4^{+/-}$ (n=7), $N4^{-/-}$ (n=5).

Fig. S9 Dermal blood vasculature was unaffected in mice with LEC loss of canonical Notch signaling**.** $Prox1CreER^{T2}$ and $DNMAML^{fl/fl}$ mice were crossed and tamoxifen administered at E12.5 and dorsal dermis analyzed at E14.5. **a**) CD31 staining of $Prox1CreER^{T2};DNMAML^{fl/+}$ ($DNMAML^{LEC}$) mutant and $DNMAML^{fl/+}$ (control) dermis. White dashed line denotes the midline. Scale bars, 1000µm. **b**) Quantification of average CD31 intensity normalized by area. Data presented ± s.e.m. control (n=3), $DNMAML^{LEC}$ (n=6).

Muley/Kim et al. Supp. Figure 1

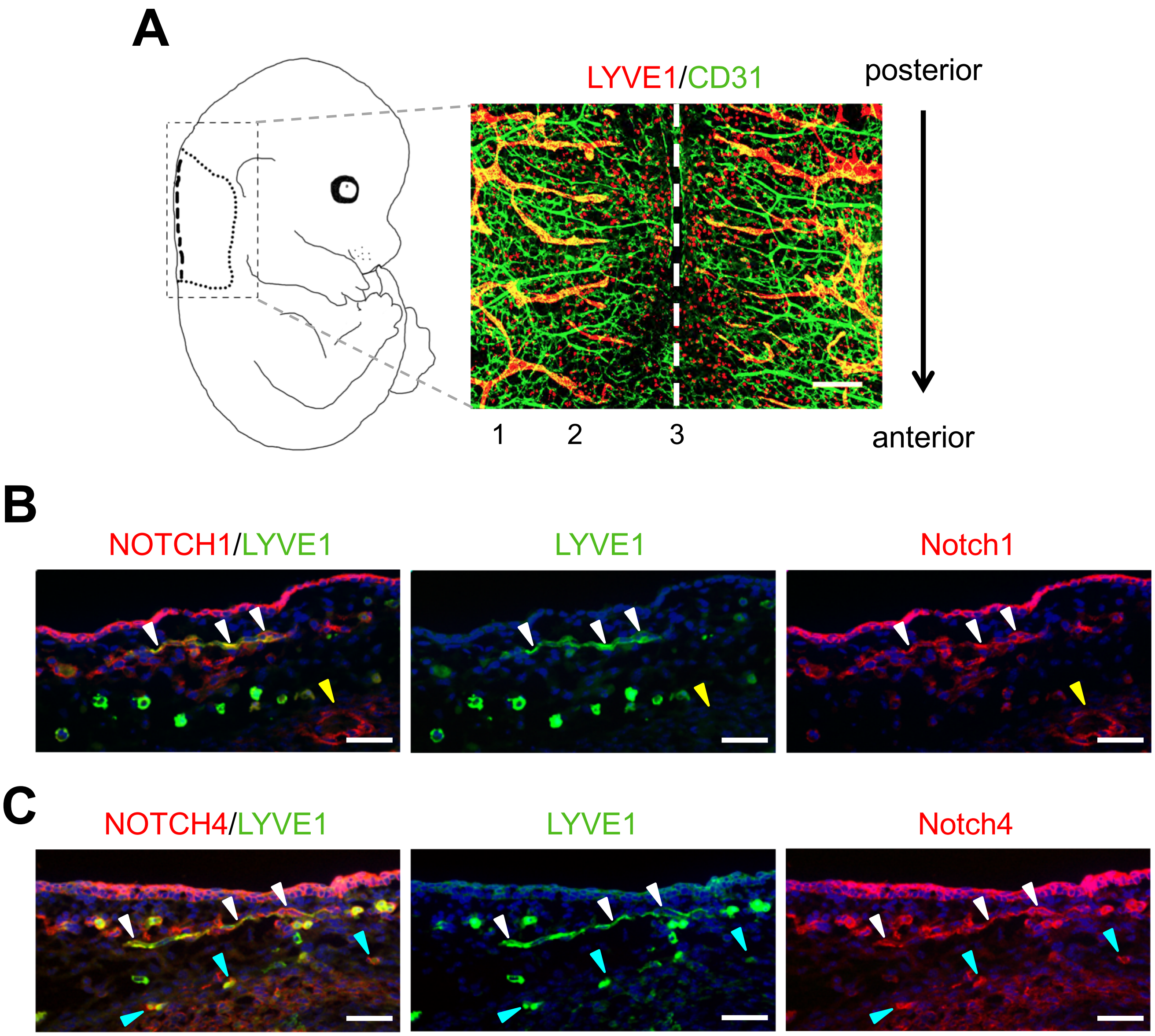


Fig. S1 Murine dermal lymphangiogenesis and NOTCH1 and NOTCH4 expression. **a**) The lymphangiogenic plexus consists of a maturing plexus (1) and active lymphangiogenic fronts (2) which migrates towards the lateral midline (3, dashed line). LYVE1 and CD31 staining of E14.5 dorsal skin. Scale bar, 100µm. **b, c**) Cross-sections of wild-type dermis stained for LYVE1 and **b**) NOTCH1 or **c**) NOTCH4. White arrowheads mark LYVE1+/NOTCH1+ and LYVE1+/NOTCH4+ LECs. Yellow arrowheads mark NOTCH1+ blood vessels. Blue arrowheads mark LYVE1+/NOTCH4+ macrophages. Scale bars, 50µm.

Muley/Kim et al., Supp. Figure 2

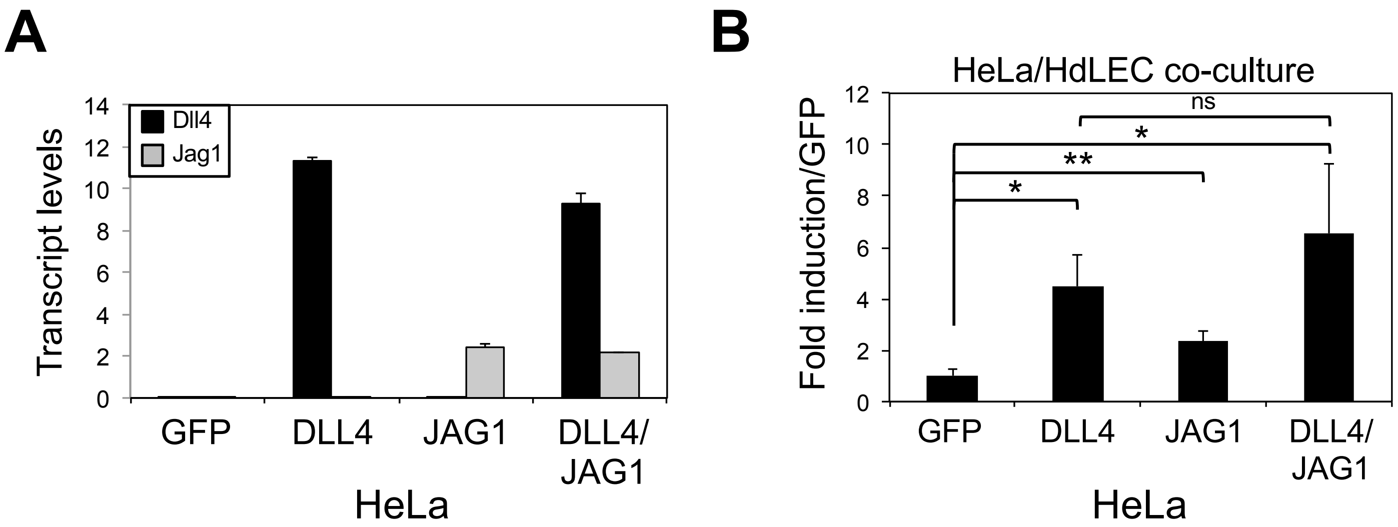


Fig. S2 Dll4 and Jag1 activated Notch signaling in HdLECs. **a**) Quantification of ectopic *Dll4* and *Jag1* transcript levels in HeLa cells expressing GFP, DLL4, JAG1 or both DLL4 and JAG1. *Dll4* and *Jag1* qRT-PCR of HeLa cell lines used in co-culture. Data presented as relative transcript levels ± s.e.m. **b**) Notch/CSL luciferase reporter assay of ligand-expressing or GFP-expressing HeLa cells (y-axis) co-cultured with HdLEC. Data presented as fold luciferase induction over HdLEC co-cultured with GFP-HeLa controls ± s.e.m. Representative experiments presented and performed in triplicate. * $p < 0.02$, ** $p < 0.001$. ns – not significant.

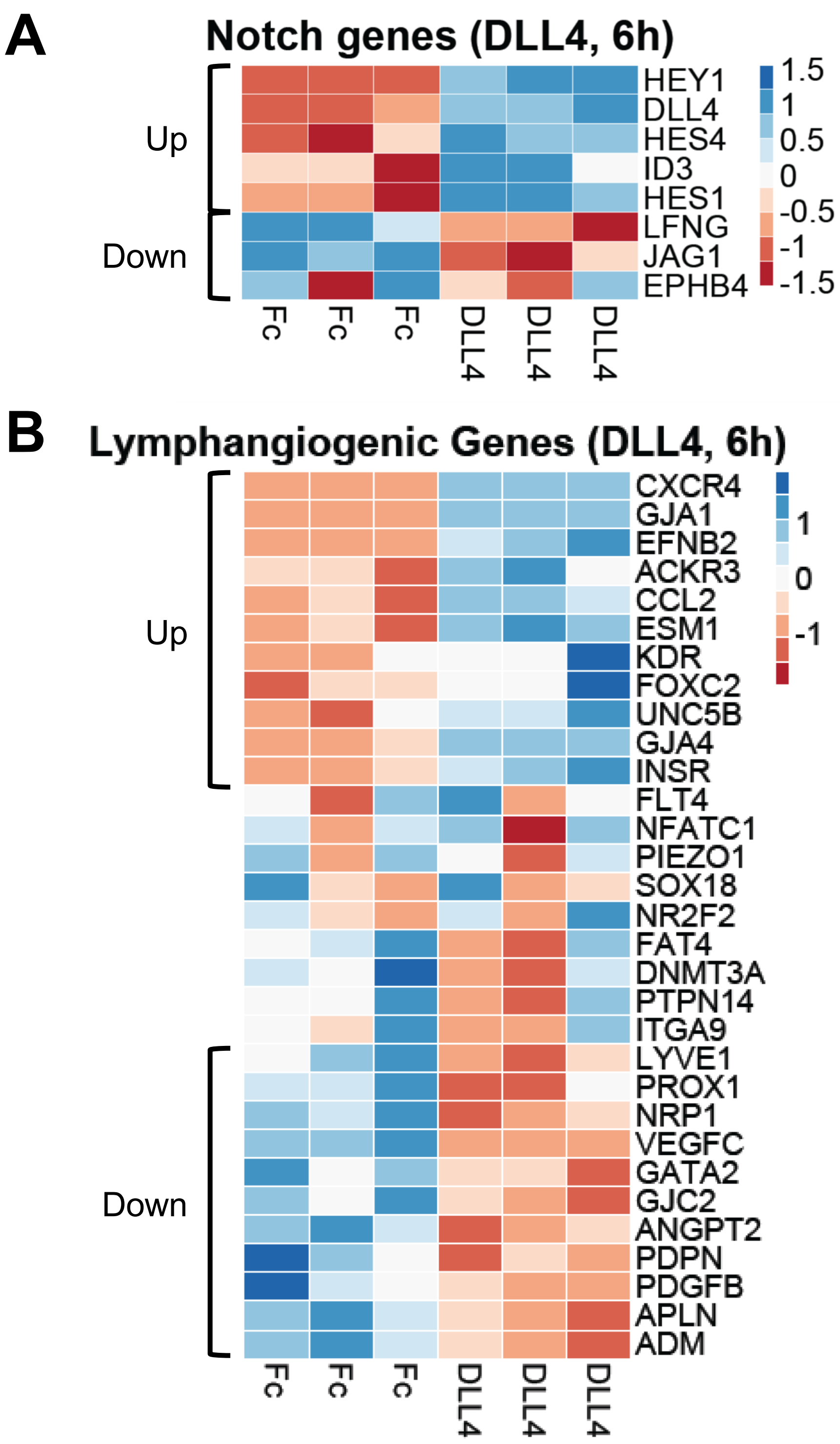


Fig. S3 Effect of DLL4/Notch signaling on Notch pathway and lymphangiogenic genes in HdLECs. **a**) Heatmap of Notch pathway genes. **b**) Heatmap of lymphangiogenic genes.

A

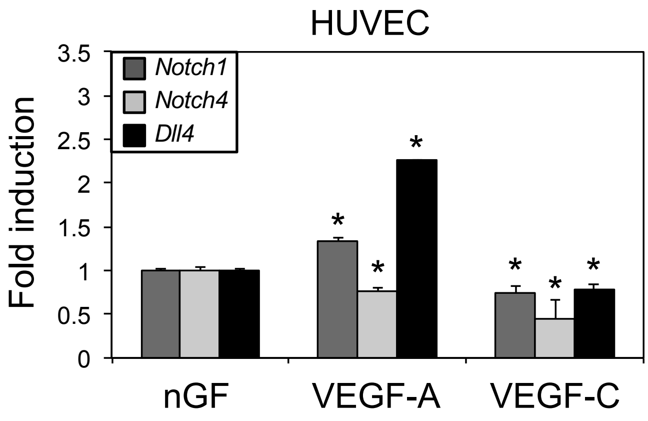


B

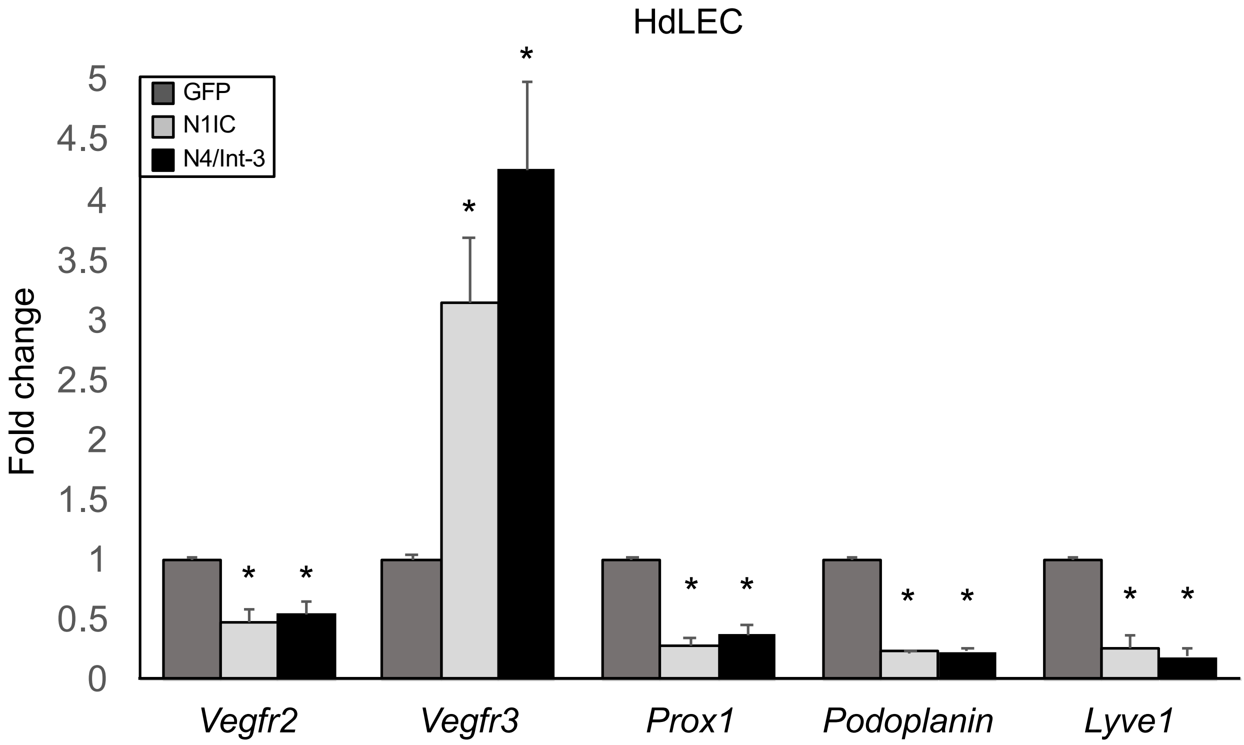


Fig. S4 VEGF and Notch signaling in HUVEC and HdLECs. **a**) *Notch1*, *Notch4*, and *Dll4* qRT-PCR of HUVEC treated for 5 hours with either VEGF-A or VEGF-C. Data presented as fold induction over HUVEC in no growth factors ± s.d. * p< 0.05. b) *Vegfr2*, *Vegfr3*, *Prox1*, *Podoplanin*, and *Lyve1* qRT-PCR of HdLEC expressing GFP, N1IC, or N4/Int-3. Data presented as fold induction over GFP expressing HdLECs ± s.e.m. Mean for > three experiments presented. T-Test * p< 0.05.

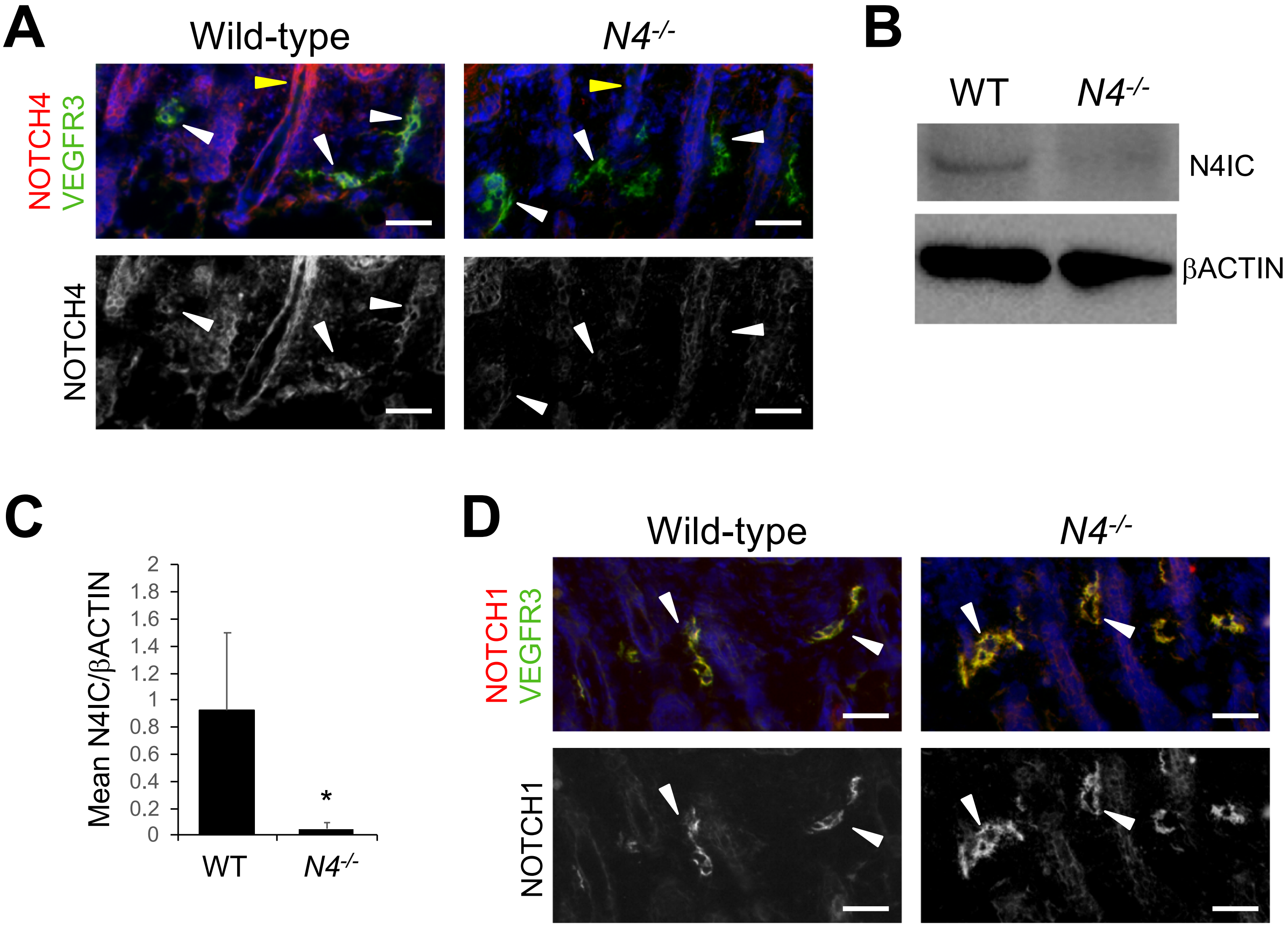


Fig. S5 Validation of the *Notch4* null mice. **a**) P4 dorsal dermal tissue cross-sections from *Notch4*$^{-/-}$ and wild-type littermates were stained for VEGFR3 and NOTCH4. NOTCH4 expression was absent in the lymphatic endothelium (white arrowheads) and epithelial cells of the hair follicle (yellow arrowheads) and dermis in *Notch4*$^{-/-}$ tissues. **b**) Representative Western blot of E14.5 *Notch4*$^{-/-}$ and wild-type tissues probed with antibodies against the cytoplasmic domain of NOTCH4 (N4IC) or βACTIN. **c)** Mean N4IC expression normalized by βACTIN for wild-type (n=3) and *Notch4*$^{-/-}$ (n=4) Westerns determined by densitometry. Data presented ± s.e.m. T-test: * $p < 0.05$. **d**) P4 dorsal dermal tissue cross-sections from *Notch4*$^{-/-}$ and wild-type littermates were stained for VEGFR3 and NOTCH1. NOTCH1 expression was unchanged in the VEGFR3+ lymphatics (white arrowheads). Scale bars, 50µm.

Muley/Kim et al. Supp. Figure 6

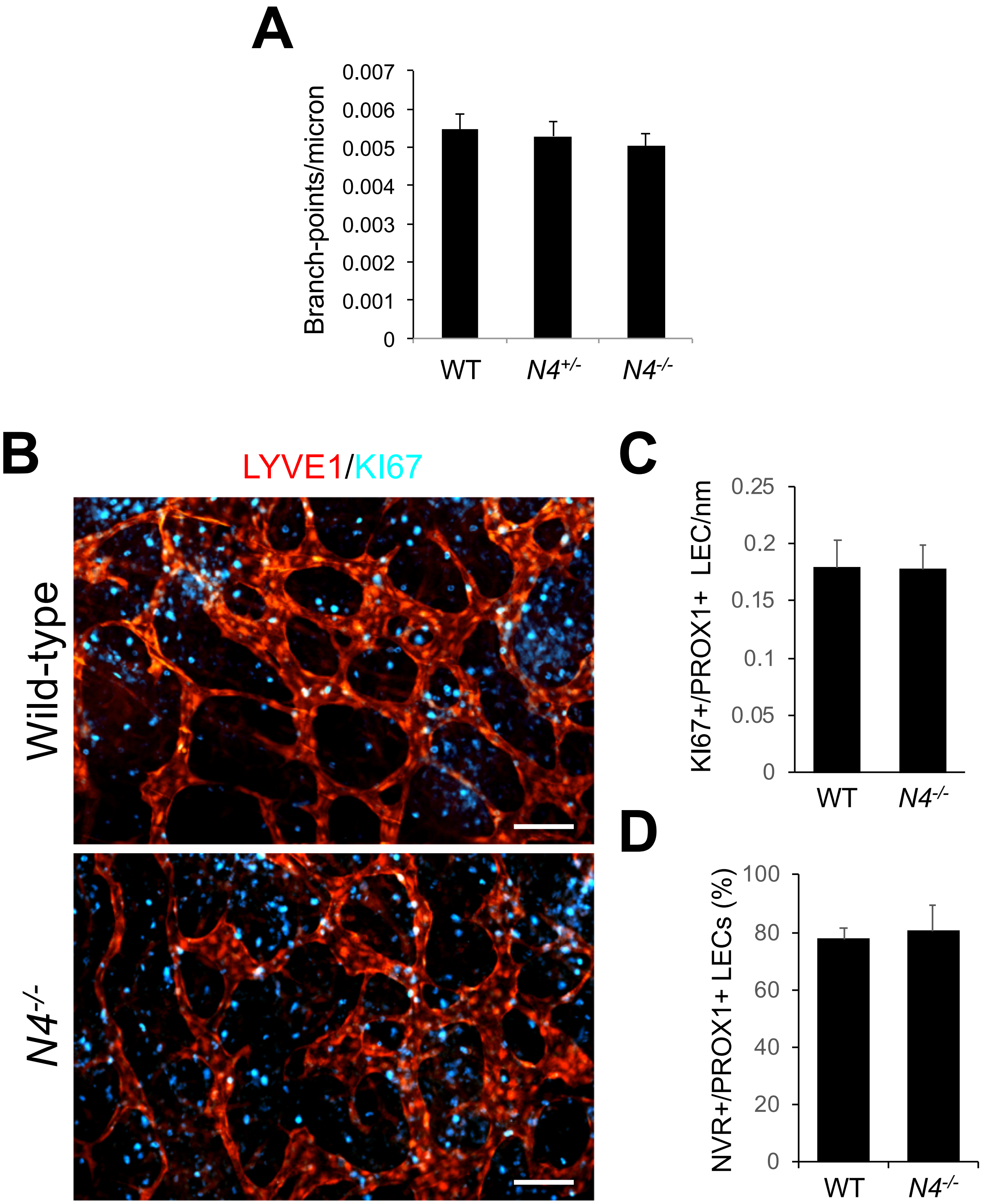


Fig. S6 Lymphatic branching, proliferation and canonical Notch signaling was unaffected in E14.5 *Notch4*$^{-/-}$ dermis. **a)** Quantification of the average number of branch-points normalized to unit of vessel length. Data presented ± sem. wt (n=7), *N4*$^{+/-}$ (n=13), *N4*$^{-/-}$ (n=7). **b**) LYVE1 and KI67 staining of E14.5 *Notch4*$^{-/-}$ and wild-type dermis. Scale bars, 100μm. **c**) Quantification of PROX1+/KI67+ LECs normalized to lymphatic vascular area ± s.e.m. wt (n=3), *N4*$^{-/-}$ (n=4) **d**) Number of PROX1+ and PROX1+/NVR+ LECs was determined for individual sprout at the lymphangiogenic front until the first branch-point for E14.5 *ProxTOM;NVR* and *Notch4*$^{-/-}$*;ProxTOM;NVR* wholemounts stained for LYVE1, Data presented as percent PROX1+/NVR+ LECs relative to total Prox1+ LECs ± s.e.m. wt (n=6), *N4*$^{-/-}$ (n=3)

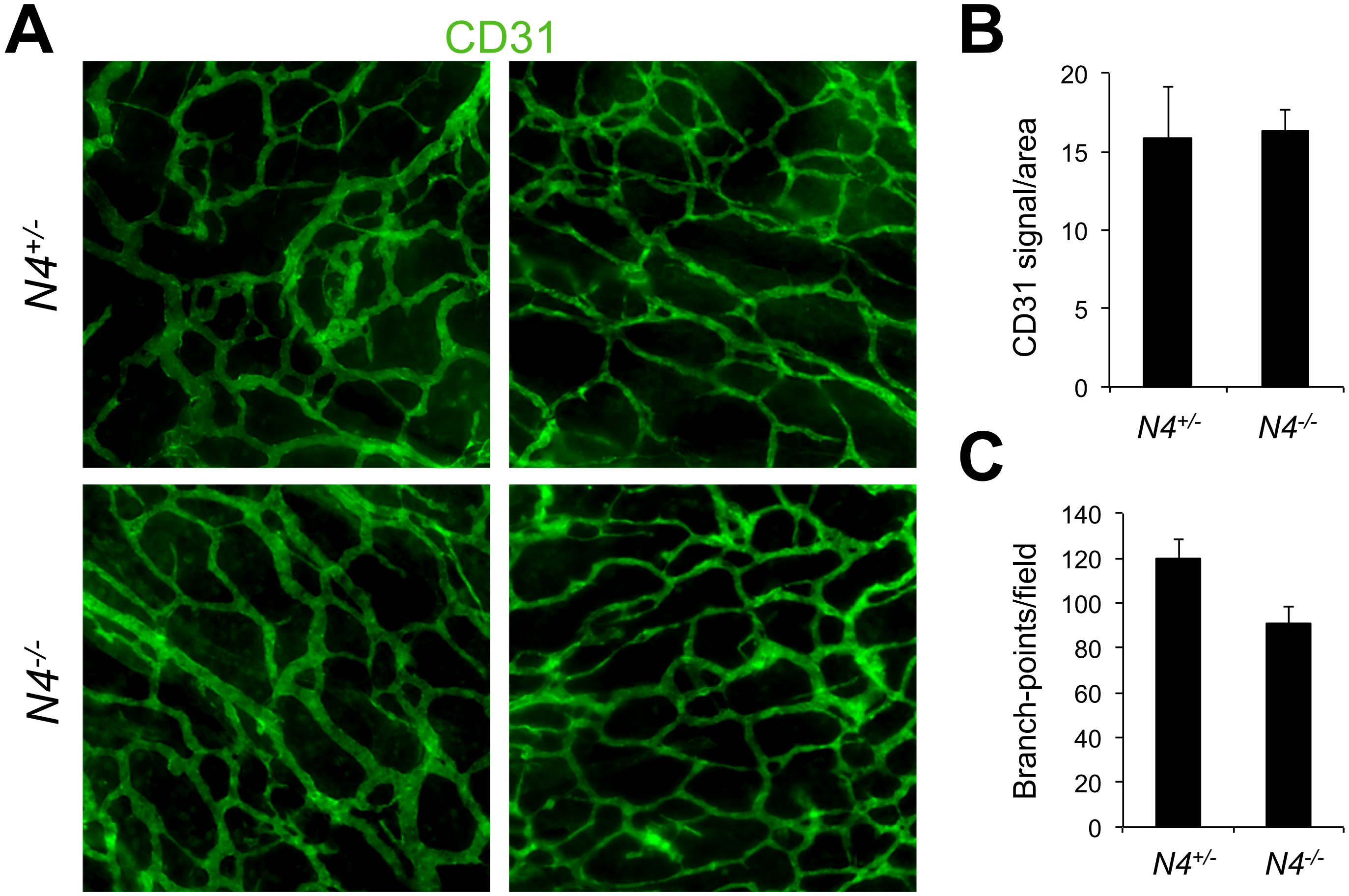


Fig. S7 Dermal blood vasculature was unaffected in *Notch4*$^{-/-}$ mice. **a**) CD31 staining of E14.5 dorsal dermal wholemounts from *N4*$^{-/-}$ and control *N4*$^{+/-}$ littermates. **b**) Quantification of average CD31 intensity normalized by area. Data presented ± s.e.m. *N4*$^{+/-}$ (n=3), *N4*$^{-/-}$ (n=4). **c**) Quantification of the average number of branch-points per field of view. Data presented ± sem. *N4*$^{+/-}$ (n=3), *N4*$^{-/-}$ (n=5).

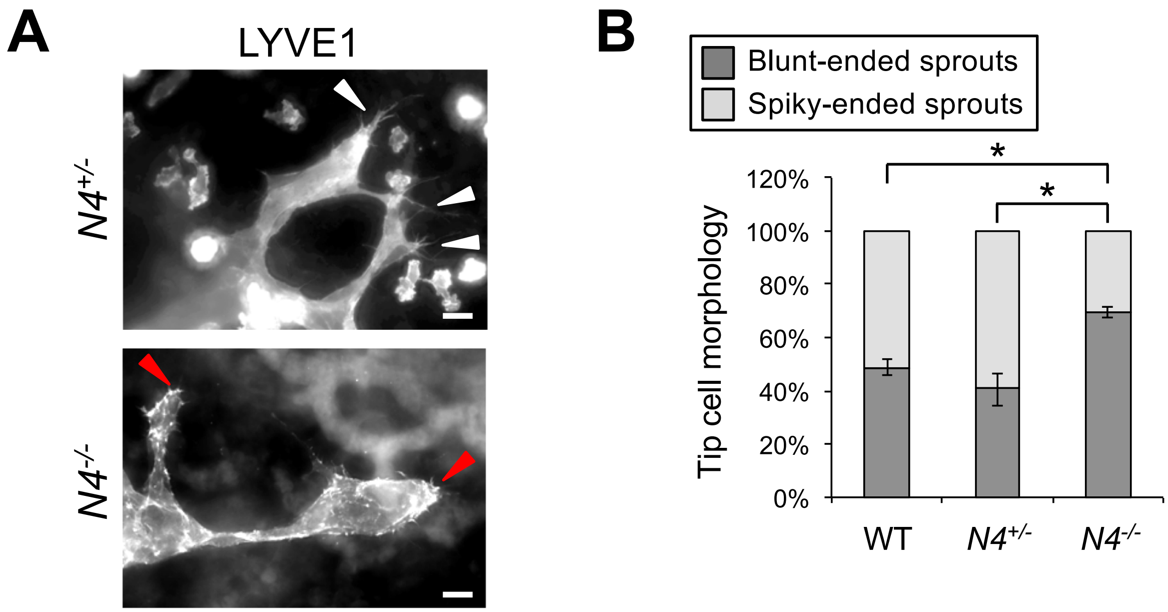


Fig. S8 Increase in blunt-ended lymphatic sprouts in E14.5 *Notch4*$^{-/-}$ dermis. **a)** High magnification images of LYVE1+ sprouts at the lymphangiogenic front. White arrowheads mark filopodia extending from a spiky-ended sprouts. Red arrowheads mark blunt-ended sprouts, Scale bars, 20µm. **b)** Quantification of the sprout morphology at the lymphangiogenic front. Blunt-ended sprouts are rounded with reduced filopodia. Spiky-ended sprouts have multiple filopodia. Data presented as percent blunt or spiky phenotype relative to total sprouts ± s.e.m. One-way ANOVA: $p= 0.004$, T-Test: $*p< 0.003$, wt (n=5), *N4*$^{+/-}$ (n=7), *N4*$^{-/-}$ (n=5).

Muley/Kim et al., Supp. Figure 9

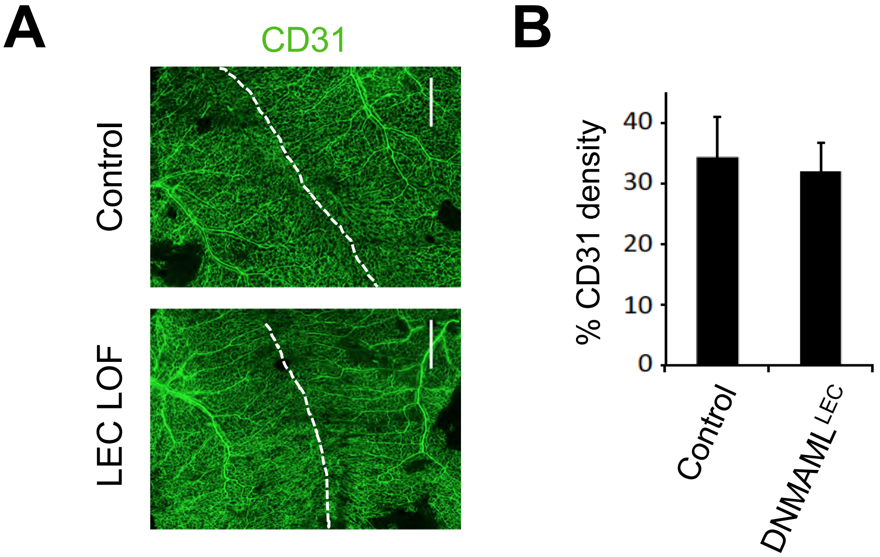


Fig. S9 Dermal blood vasculature was unaffected in mice with LEC loss of canonical Notch signaling**.** *Prox1CreER*$^{T2}$ and *DNMAML*$^{fl/fl}$ mice were crossed and tamoxifen administered at E12.5 and dorsal dermis analyzed at E14.5. **a**) CD31 staining of *Prox1CreER*$^{T2}$*;DNMAML*$^{fl/+}$ (*DNMAML*$^{LEC}$) mutant and *DNMAML*$^{fl/+}$ (control) dermis. White dashed line denotes the midline. Scale bars, 1000µm. **b**) Quantification of average CD31 intensity normalized by area. Data presented ± s.e.m. control (n=3), *DNMAML*$^{LEC}$ (n=6).
